# A prioritized medium-throughput screen identifies fibroblast growth factors as protective for human stem cell derived insulin secreting beta cells

**DOI:** 10.1101/2025.02.12.637990

**Authors:** Chieh Min Jamie Chu, Sepehr Kamal, Luo Ting Huang, Carmen L. Bayly, Maria J. Beletsky, Samantha Mar, Sing-Young Chen, Meltem E. Omur, Wyeth W. Wasserman, Francis C. Lynn, James D. Johnson

## Abstract

Stem cell-derived β-like cells (SCβ cells) are an alternative to cadaveric β cells for replacement therapy in type 1 diabetes. However, SCβ cells face a multitude of stresses. Both β cells and SCβ cells reside in complex microtissues and are modulated by hundreds of autocrine/paracrine signals. Here, we leveraged multi-omics data from late-stage SCβ cells and human islets to map ligand-receptor pairs and generate a prioritized list of ligands for high-content SCβ cell survival screening. Our medium-throughput screen tracked cell number, cell death, and *INS* production over several days using automated, high-content imaging. Members of the fibroblast growth factor (FGF) family significantly prevented cytokine-induced cell death, with the top validated hits being FGF4, FGF5, and FGF8F. With these results, there is the potential to improve SCβ cell survival *in vitro* via readily targetable pathways and to produce a more robust product for translation to the clinic.

## Introduction

Type 1 diabetes is the result of autoimmune destruction of pancreatic β cells, which are responsible for maintaining glucose homeostasis via production of the hormone ligand insulin. Cell replacement therapies with stem cell-derived β cells (SCβ cells) have been proposed to address this loss of β cells(Hebrok 2012, Kolic and Johnson 2021). Several issues need to be addressed before SCβ cells are ready for broad clinical use. For example, current SCβ cells do not exactly match the insulin production and secretion patterns of primary human β cells(Hebrok 2012, Kolic and Johnson 2021, Maghera, Ellis et al. 2026). Also, the resilience of SCβ cells during their differentiation, transplantation, engraftment and function in autoimmune and alloimmune environments remains a challenge(Atkinson, Roep et al. 2019). Previous studies have endeavoured to find methods and factors that promote β cell survival using CRISPR, chemical, and shRNA screens(Cai, Ishikawa et al. 2020, Sintov, Nikolskiy et al. 2022, Rebecca A Lee 2024, Vandana, Zhu et al. 2025).

Pancreatic β cells, are subjected to many stresses, including the production of insulin itself(Szabat, Page et al. 2016). Presumably, the same stresses may impact SCβ cell fragility and, if unmanaged, could result in SCβ cell death and dysfunction(Brozzi and Eizirik 2016, Mirmira, Sims et al. 2016). While studies have been done to understand the effects of various sources of stress in SCβ cells, most lack temporal information and do not provide information on dose effects(Demine, Schiavo et al. 2020, Castro-Gutierrez, Alkanani et al. 2021, Dettmer, Niwolik et al. 2022). Therefore, it is important to comprehensively examine how SCβ cells respond to stress in a time and dose dependent manner, and to find ways to alleviate these stresses to prevent excessive SCβ cell loss and to reduce the number of cells needed per transplant.

Autocrine and paracrine signaling processes are critical for the differentiation and function SCβ cells, but they remain poorly characterized. In fact, cell surface receptors are the target of 90% of small molecules and ligands used in SCβ cell differentiation protocols and these signaling pathways are thought to at least partially mimic signalling that occurs during endocrine cell development(Rezania, Bruin et al. 2014, Russ, Parent et al. 2015). With advances in omics technologies and public datasets, it should be possible to elucidate missing factors and reveal novel pathways that could be leveraged to improve SCβ cell survival.

Efforts to identify the protein ligands, receptors, and interacting ligand-receptor pairs in the human genome included the Database of Ligand-Receptor Partners (DLRP), IUPHAR, and Human Plasma Membrane Receptome (HPMR)(Graeber and Eisenberg 2001, Ben-Shlomo, Yu Hsu et al. 2003, Sharman, Benson et al. 2013). Further efforts to identify protein interactions through publicly available data include revisions on these lists in 2015, connectomeDB2020 in 2020, and OmniPath in 2021(Franceschini, Szklarczyk et al. 2013, Ramilowski, Goldberg et al. 2015, Hou, Denisenko et al. 2020, Turei, Valdeolivas et al. 2021). These datasets and analyses have greatly increased our understanding of cell-to-cell signaling and the ligands and proteins involved. There is an unmet need for comprehensive and curated analyses of intra-islet signaling that compares SCβ cells and human islets. We previously generated a list of factors and ligands which were used to screen for alleviators of cell death in mouse primary β cells(Yang, Wills et al. 2015) and here we sought to extend this approach to human islets and stem cell-derived islet-like clusters. In the present study, we generate a prioritized list of ligand-receptor pairs from bulk RNAseq, single cell RNAseq (scRNAseq), and proteomics datasets. We screened these prioritized ligands for their ability to protect SCβ cells in the context of cytokine stress. Our screen revealed fibroblast growth factors (FGFs) as ligands that promote SCβ cell survival and function.

## Results

### Curation of islet and stem cell-derived islet ligand and receptor resources

Ligand-receptor databases contain many entries that are likely not true ligands or receptors due to a combination of incorrect functional or locational annotations(Kunin and Ouzounis 2005, Schnoes, Brown et al. 2009, Promponas, Iliopoulos et al. 2015) and/or excessively broad definitions of the terms “ligand” and “receptor”. For this study, we focused on the subset of cell-cell signaling involving secreted protein ligands and the plasma membrane receptors that have literature-supported interactions. We excluded intracellular receptors, ECM proteins, major histocompatibility complex proteins, non-protein ligands, complement proteins, protein complexes, and extracellular enzymes from the initial list of 927 protein ligands and 4285 receptors from OmniPath(Turei, Korcsmaros et al. 2016). We judged that many of the listed ‘receptors’ were not likely to be cell surface receptors for protein ligands that met our criteria. The OmniPath consensus score measures the number of resources that include each ligand or receptor. We used the OmniPath consensus score as a measure of literature support to select only ligands and receptors whose roles are well established. We included only ligands with a consensus score above four and only receptors with a consensus score above seven (Supplemental Figure 1A). This excluded 3698 receptors with a consensus score of one (Supplemental Figure 2A). We then manually reviewed the remaining 490 ligands and 587 receptors; we excluded an additional 68 ligands and 238 receptors (Supplemental Figure 2B).

Examples of ‘ligands’ that we manually excluded were the ECM protein LAMA1, the complement protein C4B, and the membrane-bound ligand CD40LG. Examples of receptors that we manually excluded were the ECM protein ITGB2 and the non-protein ligand receptor ADORA1. After manual filtering, we had a final list of 422 protein ligands and 349 receptors. The top ligands based on OmniPath consensus score were BMP2, IL10, and INHBA, and the top receptors were KDR, EGFR, and TGFBR1. The signaling pathways with the highest ligand count in our final list were CCL, WNT, and FGF, and the signaling pathways with the highest receptor count were ncWNT, CCL, and EPHA (Supplemental Figure 2C). Of the top represented pathways, the IL4, NT, and IL2 pathways had the highest mean consensus scores for the ligands, and the IL4, CXCL, and NOTCH pathways had the highest mean consensus scores for the receptors (Supplemental Figure 2C). These pathways have robust literature support for their functions in intercellular signaling.

We next identified the interacting pairs between our curated ligand and receptor lists. We obtained an initial list of 40,014 ligand-receptor interaction pairs with literature support from OmniPath. After filtering to include only the selected 422 ligands and 349 receptors, we were left with 1552 interaction pairs (Figure 1A). 26 ligands and 27 receptors had no interaction pairs. The mean number of interactions was 3.68 per ligand and 4.45 per receptor. The ligands with the most interactions were POMC and WNT5A, with 14 interactions each. Interestingly, the receptors with the most interactions had much higher interaction counts than the ligands (Figure 1B). The receptors with the most interactions were MTNR1A, ACVR2B, and ACVR2A, with 44, 27, and 26 interactions, respectively.

**Figure 1.**
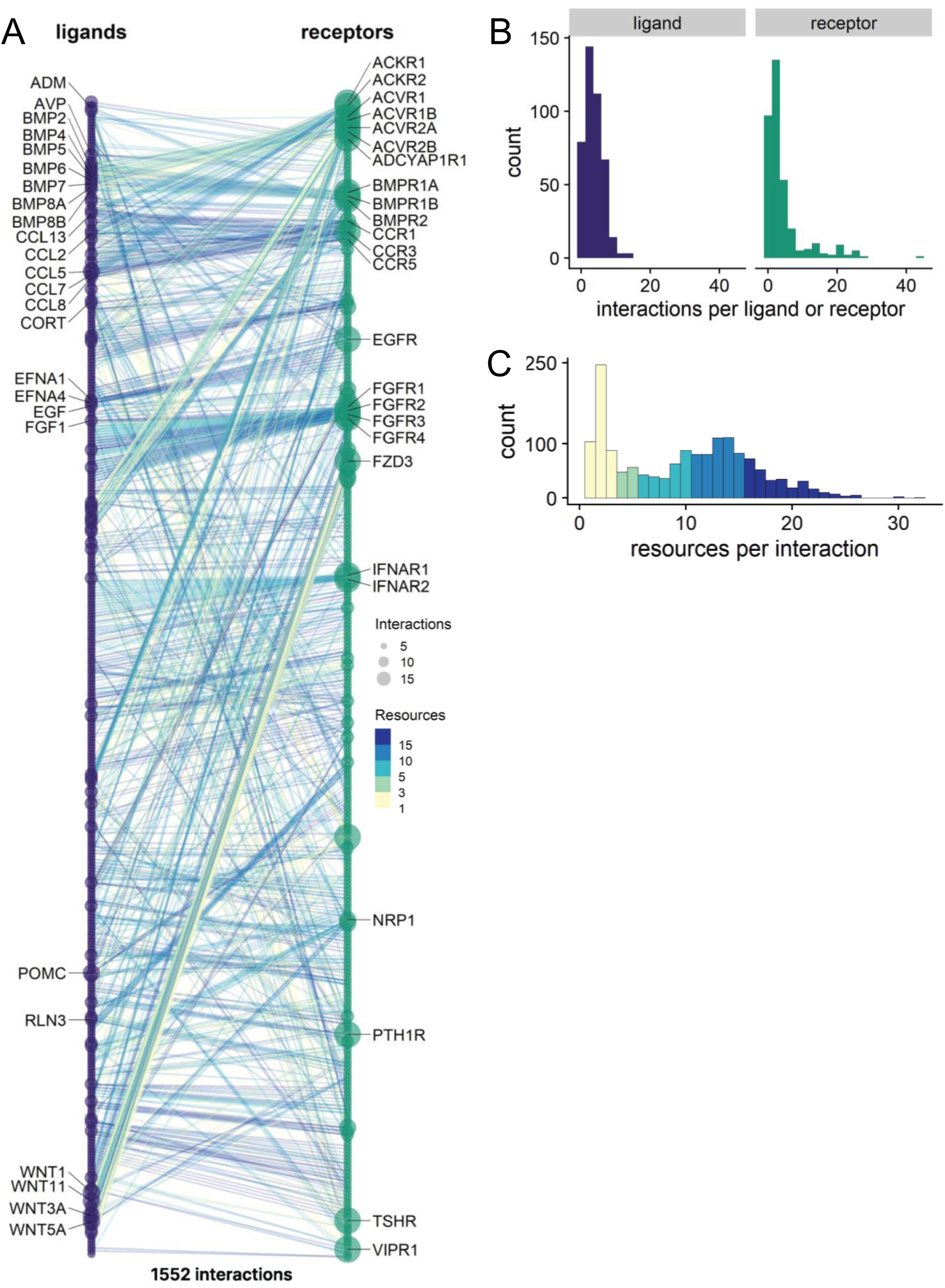
Curation of ligand-receptor pairs and interactions. **(A)** All 1552 interactions between the final filtered list of ligands (left) and receptors (right) are shown in alphabetical order. 26 out of 422 ligands and 27 out of 349 receptors had no interactions and are not shown. Each line represents a ligand-receptor pair. The line color indicates the number of resources supporting the interaction. The dot size indicates the number of interactions per ligand or receptor. Labels indicate the top 25 ligands and receptors with the most interactions, with at least 5 resources supporting their interactions. **(B)** Distribution of the number of interactions for each ligand and receptor. **(C)** Distribution of the number of resources in OmniPath that support each of the 1552 interactions. **(D)** Overview of the ligand-receptor interactions in our custom resource.

The number of resources which supported each interaction ranged from 1 to 32 (Figure 1C). The interactions with the most literature support were TNF-TNFRSF1A, EGF-EGFR, TNF-TNFRSF1B, EPO-EPOR, and HGF-MET. Our curated ligand-receptor resource can be used by us and others to analyze cell-cell signaling in islets and create prioritized lists for medium-throughput screens.

### Correlation of gene expression and protein abundance in human islets

While transcriptomes typically have deeper coverage than proteomes, proteins are the primary actuators of cellular physiology and signaling. Therefore, we incorporated RNAseq data and proteomics data from the same 90 human islet donors(Kolic, Sun et al. 2024)(MacDonald-Gloyn 2023) to provide orthogonal data on ligand and receptor proteins. Of the 8194 proteins detected in the proteomics, there were 41 ligands and 41 receptors. It should be noted that protein abundances cannot be directly compared between gene products due to their differential appearance in lysates, sensitivity to enzymatic digestion, and detection on mass-spectroscopy. We also analyzed data from another dataset (n=89) that was not donor matched from Lund 2014. We determined the mRNA-protein correlation using both the Lund 2014 and donor-matched MacDonald-Gloyn 2023 datasets.

Interestingly, the donor-matched RNA-seq dataset did not have a stronger protein-mRNA correlation than the non-donor-matched Lund 2014 dataset (Figure 2A,B). Ligands (R = 0.55, *p* value < 0.0002) had a significantly higher protein-mRNA correlation than the receptors (R = 0.02, *p* value < 0.91). The correlation for the ligands was slightly higher than the genome-wide correlation of R = 0.49 (Figure 2A,B). These results indicate protein abundance did not correlate with mRNA abundance for islet cell receptors. The ligands with the lowest protein to RNA ratio were IL16, BMP1, and ANGPTL4, and the receptors with the lowest protein to RNA ratio were PLXNA1, EPHB4, and IL3RA (Figure 2A,B). Overall, we found a correlation between protein abundance and gene expression for ligands and a relative lack of correlation for receptors in human islets.

**Figure 2.**
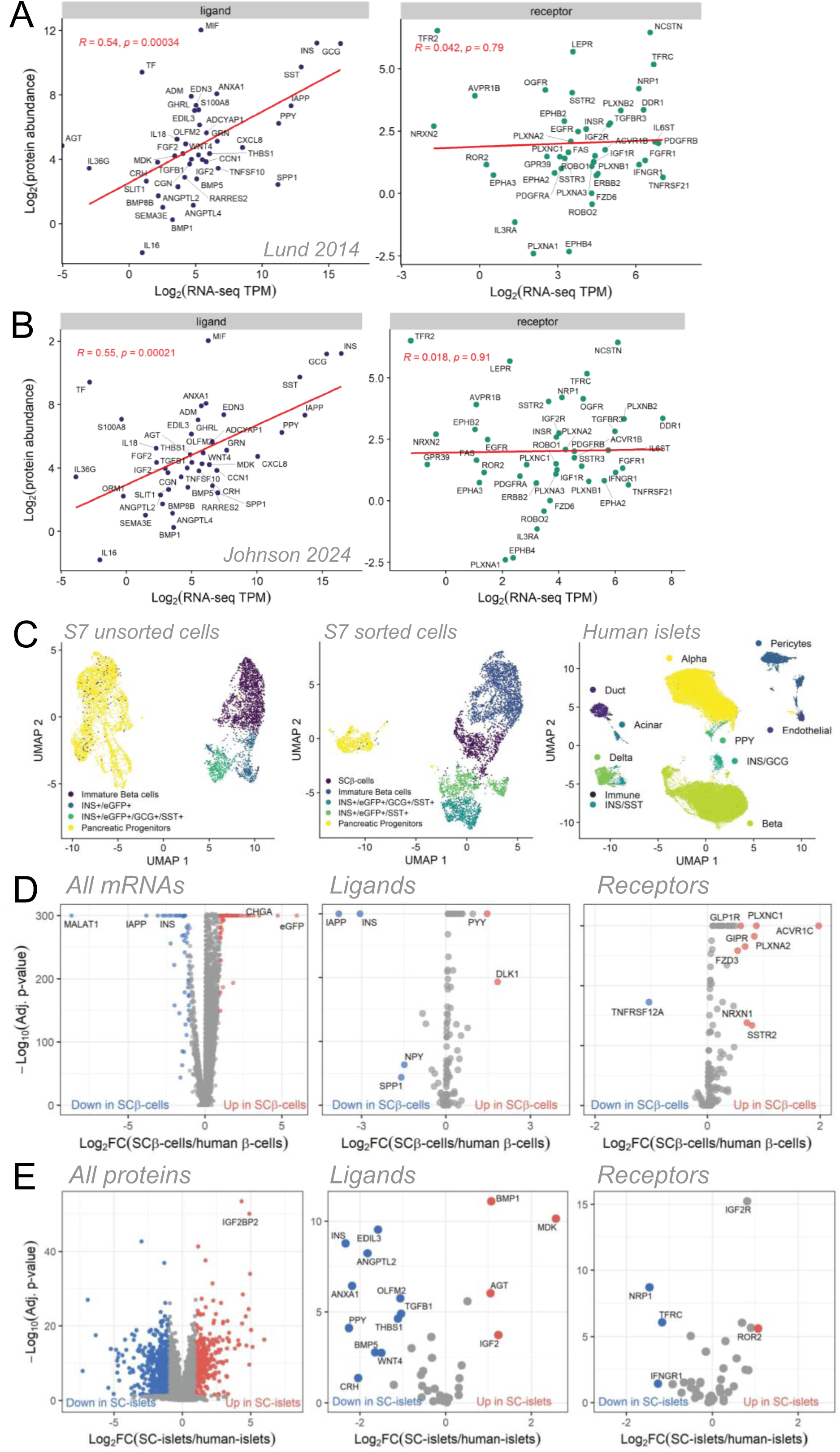
Analyses on gene expression and protein abundance from bulk RNAseq, scRNAseq, and proteomics datasets. **(A,B)** Correlation between proteomics (n=90) and two separate bulk RNA-seq datasets for all ligands (left) and receptors (right) detected in the proteomics. Protein abundances were normalized to the amino acid length of the proteins. **(A)** Lund 2014 bulk RNA-seq dataset (n=89). **(B)** MacDonald-Gloyn 2023 bulk RNA-seq dataset (n=90) with donor-matched samples to the proteomics. **(C)** UMAP projections for S7 unsorted SC-islet (n=6875 cells), S7 sorted SC-islet (n=5355 cells), and human islet (n=68,650 cells) scRNA-seq datasets. **(D)** Differential gene expression between SCβ cells (n=891 cells) and human β cells (n=29,956 cells) for all genes (left), ligands (middle), and receptors (right). Genes in red and blue are significantly differentially expressed (absolute log2FC > 1 and adjusted *p* value < 0.05). Genes just below the absolute fold change threshold but with significant *p* value are also labeled. **(E)** Differential protein abundance between non-diabetic human islets (n=118) and SC-islets (n=18). Proteins in red and blue are significantly differentially abundant (absolute log2FC > 1 and adjusted *p* value < 0.05). All 8194 proteins detected in proteomics (left). IGF2BP2 is highlighted as it regulates IGF2 expression. Differentially abundant ligands (middle) and receptors (right).

*Differential gene expression between SC*β *cells and human* β *cells*

We next compared ligand and receptor expression in scRNA-seq data of human islets, stage 7 unsorted SC-islets, and stage 7 sorted SC-islets. Specifically, we selected the SCβ cell cluster from the stage 7 sorted SCβ cells to compare to human β cells (Figure 2C). The clusters contained 891 SCβ cells and 29,956 human β cells (Figure 2C). As expected for this knock-in cell line, we found EGFP was the top elevated gene in SCβ cells (Figure 2D). We found two ligands increased and four decreased in SCβ cells compared to human β cells (Figure 2D). The increased ligands were *DLK1* and *PYY*, and the decreased ligands were *IAPP*, *INS*, *SPP1*, and *NPY*. Interestingly, *IAPP* and *INS* are canonical β cells maturity markers(Muraro, Dharmadhikari et al. 2016, Bayly, Dai et al. 2024), while *NPY* is a marker of immature β cells(Rodnoi, Rajkumar et al. 2017). One receptor was significantly higher (*ACVR1C*) and one was significantly lower (*TNFRSF12A*) in SCβ cells compared to human β cells (Figure 2D); however, multiple additional receptors were just below significance thresholds (Figure 2D). *INS*, *ACVR1C*, and *TNFRSF12A* expression were evenly distributed among cells in the SCβ cell and human β cell clusters, however *IAPP* expression was limited to only a subset of the SCβ cells (Supplemental Figure 3A). This subset may correspond to a more mature subpopulation of SCβ cells(Bayly, Dai et al. 2024). This analysis identified the top differentially expressed ligands and receptors between SCβ cells and human β cells.

### Differential protein abundance between human islets and SC-islets

We compared ligand and receptor protein abundances between non-diabetic human islet donors and a mixed pool of S7 sorted and S7 unsorted *INS*-EGFP SC-islets. We found 4 ligands higher and 11 lower in SC-islets compared to human islets (Figure 2E). The top increased ligands were MDK, BMP1, and IGF2, and the top decreased ligands were INS, ANXA1, and PPY (Figure 2E). There was 1 receptor higher (ROR2) and 3 receptors lower (NRP1, TFRC, IFNGR1) in SC-islets compared to human islets (Figure 2E). While its fold change was just below the threshold for inclusion, IGF2R was also significantly higher in SC-islets. Interestingly, IGF2BP2, which binds to and regulates the translation of IGF2 mRNA(Nielsen, Christiansen et al. 1999, Dai, Rapley et al. 2011), was also one of the top increased proteins genome-wide (Figure 2E). The combination of IGF2, IGF2R, and IGF2BP2 elevation makes this the top upregulated signaling pathway at the protein level in SC-islets.

### Intra-islet signaling pathways in SC-islets and human islets

We used CellChat to expand our single-cell analysis to identify incoming and outgoing signaling pathways in human islets and S7 sorted SC-islets(Jin, Guerrero-Juarez et al. 2021). We identified 6557 total inferred interactions in human islets (Figure 3A); however, we identified only 42 total inferred interactions in S7 sorted SC-islets (Figure 3B,E). This difference is likely due to the SC-islet dataset containing 10x fewer cells. Compared to other intercellular communication tools, such as CellPhoneDB, CellChat has been shown to infer fewer interactions but capture the strongest and likely most significant interactions(Jin, Guerrero-Juarez et al. 2021). CellChat has also been shown to be fairly robust to noisy data(Dimitrov, Turei et al. 2022), allow us to draw some insights from the SC-islets analysis despite the lack of depth in the results. We found that human islets had considerably more interactions than SC-islets (Supplemental Figure 4A). We found that α-cells were by far the strongest sources of signals in human islets, while α cells, β cells, and pericytes were the strongest receivers of signals (Figure 3A). Interestingly, pericytes contributed significantly to cell-cell signaling even though they were less abundant than α cells and β cells (Figure 2C). In SC-islets, immature endocrine cells were by far the largest overall contributors to cell-cell signaling (Figure 3B), even though they were also not the largest cell population (Figure 2C). Pancreatic progenitors were the strongest senders of signals, and pancreatic progenitors, immature β cells, and SCβ cells were the strongest receivers of signals (Figure 3A). The midkine (MK) pathway was the strongest signaling pathway in both SC-islets and human islets (Figure 3C-D). In human islets, α-cells sent signals via the MK pathway to β cells, pericytes, and α-cells themselves (Figure 3C). In SC-islets, immature endocrine cells sent signals via the MK pathway mainly to themselves (Figure 3D). Human β cells both sent and received signals via the WNT pathway, while SCβ cells only sent WNT signals. Both human β cells and SCβ cells received strong signals via the IGF pathway. In human islets, the IGF signals originated from pericytes, while in SC-islets they originated from pancreatic progenitors. In human islets, duct cells and endothelial cells sent signals to β cells and α-cells via the SPP1 and ANGPT pathways. These pathways were not detected in the SC-islets. Overall, these data expand on our earlier interaction analyses, identify the signaling pathways by which islet cells communicate with each other, and highlight the contributions of non-endocrine cell types to intra-islet signaling.

**Figure 3.**
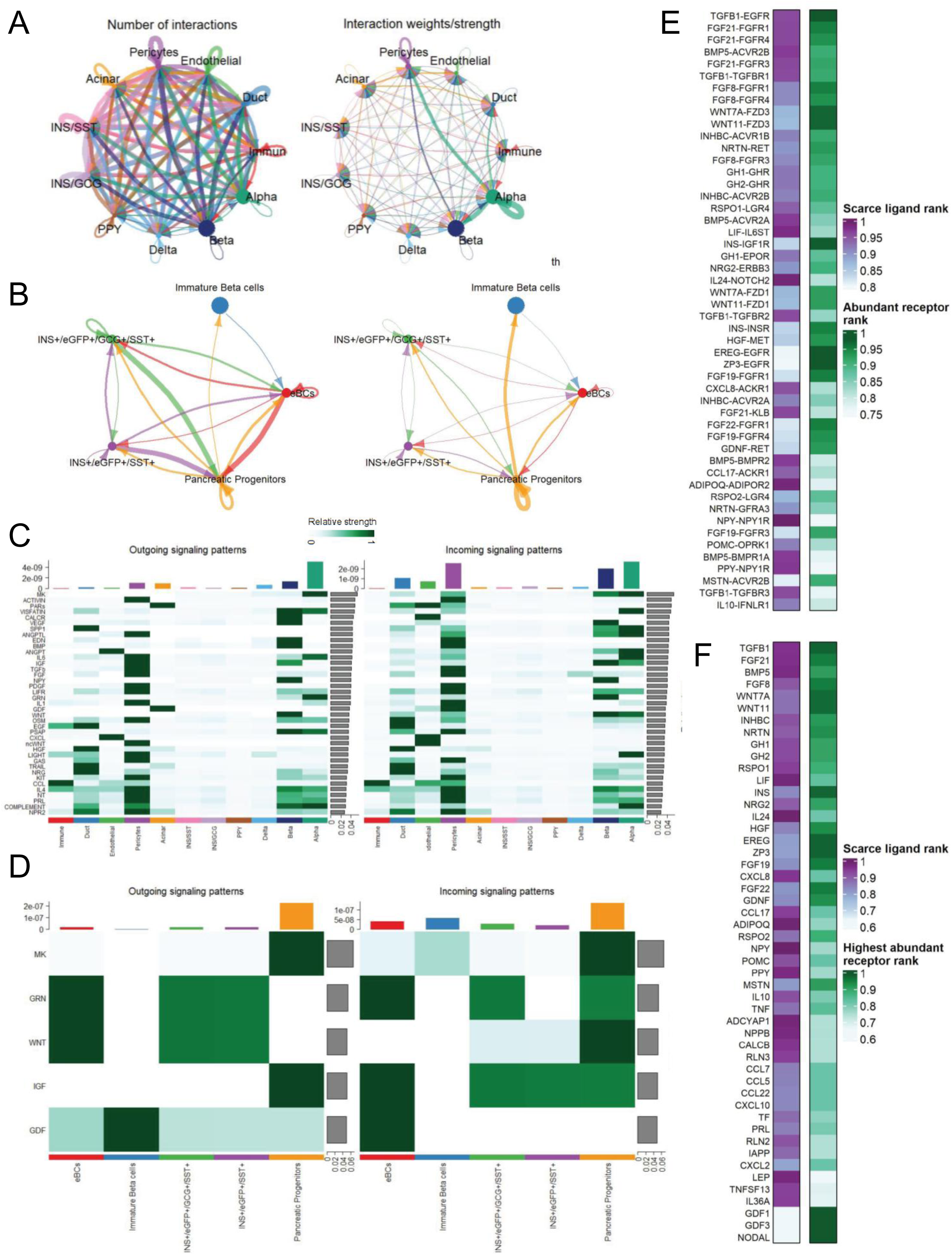
Intra-islet signaling pathways in SC-islets and human islets. **(A)** Number of interactions (left) and strength of interactions (right) between all 11 cell types in human islets. **(B)** Number of interactions (left) and strength of interactions (right) between all 5 cell types in S7 sorted SC-islets. **(C-D)** Top signaling pathways and cell types contributing to intercellular communication in **(C)** human islets and **(D)** S7 sorted SC-islets. Bar plot on top indicates the total signaling strength of each cell group. Bar plot on right hand side indicates total contribution of each signaling pathway. **(E)** Prioritized ligand-receptor interactions in SC-islets based on 3 criteria. **(F)** Top 50 scarce ligands with abundant receptors.

### Ligand and receptor prioritization in SC-islets

We combined our datasets with publicly available datasets to identify ligands and receptors in SC-islets with consistent patterns across studies, experimental methods, and differentiation protocols. We ranked ligands and receptors grouped into three categories. The three categories were expression in SC-islets, upregulation in SC-islets compared to human islets, and downregulation in SC-islets compared to human islets (Supplemental Figure 5A,B). Rankings included comparisons of bulk RNAseq, scRNAseq, and bulk proteomics from human primary islets, human primary β cells, SC-islets, and SCβ cells. We defined abundant ligands and receptors by high expression in SC-islets and higher expression in SC-islets compared to human islets. The top abundant ligands were MK, SEMA3A, and GCG (Supplemental Figure 5C). MDK was highly expressed in all five parameters and was upregulated in SC-islets in four out of five parameters. Interestingly, some ligands showed a discrepancy in differential expression between datasets. AGT was upregulated in the proteomics but was downregulated in both the Veres et al scRNA-seq and Alvarez-Dominquez et al bulk RNA-seq.

SST was upregulated in the Lynn scRNA-seq but downregulated in Veres et al and Balboa et al scRNA-seq. These discrepancies may represent biological differences between research groups in the SC-islet cells. The top abundant receptors were PLXNC1, PLXNA2, and IGF2R (Supplemental Figure 5D). They were highly expressed in all five parameters and were upregulated in four out of five parameters. ACVR1C was highly expressed and elevated in all RNA-seq datasets but was not detected in proteomics. We defined scarce ligands and receptors by low expression in SC-islets and lower expression in SC-islets compared to human islets. The top scarce ligands were EDN3, NPY, and SPP1 (Supplemental Figure 5E). These ligands had moderate expression in SC-islets and were strongly decreased in multiple parameters compared to human islets. UCN3, a marker of β cell maturity(Blum, Hrvatin et al. 2012), was decrease in SC-islets in three parameters. The top scarce receptors were TNFRSF12A, NGFR, and TNFRSF14 (Supplemental Figure 5F). TNFRSF12A was also the second highest expressed receptor in human islets based on bulk RNA-seq (Figure 2A,B) and was the only receptor lower in the Lynn scRNA-seq dataset (Figure 2D). These data identify the top ligands and receptors in SC-islets and compare their expression to human islets.

We next prioritized ligand-receptor interactions in SC-islets based on three combinations of the ligand and receptor ranks (Figure 3E,F; Supplemental Figure 6A-D. These three combinations were chosen based on the greatest differences we could identify when comparing SC-islets to primary human islets. These interactions were based on the results from our Omnipath analysis. The top interactions in SC-islets between abundant receptors and abundant ligands were SEMA3A-PLXNA2, GDF11-ACVR1C, and IGF2-IGF2R (Supplemental Figure 6A). The high rank of IGF2-IGF2R showed that the RNA-seq datasets were aligned with the proteomics, which had identified IGF2-IGF2R as the top upregulated interaction (Figure 2E). The top interactions in SC-islets with scarce receptors and abundant ligands were VEGFB-FLT1, GRN-TNFRSF1B, and NTS-NGFR (Supplemental Figure 6B). The top interactions with scarce ligands and abundant receptors were TGFB1-EGFR, FGF21-FGFR1, and FGF21-FGFR4 (Figure 3E). In all three prioritized interaction lists, some ligands and receptors appeared multiple times with different interaction partners. For example, FGFR1 was present in four of the top 50 interactions between abundant receptors and abundant ligands, and FGF21 was present in three of the top five interactions between scarce ligands and abundant receptors. To guide future screens, we simplified the three prioritized lists by selecting only the highest-ranked interaction partner for each ligand or receptor (Figure 3F; Supplemental Figure 6C, D). In summary, we prioritized the ligands, receptors, and interactions in islets by combining multiple datasets. These data improve our understanding of islet intercellular communication and enable researchers to focus future studies on ligands and receptors most likely to have a biological effect in islets.

*Live imaging of SC*β *cell response to various stressors*

We have previously used long-term, live-cell, multi-parameter, robotic imaging as a tool for tracking cell death over time**(Yang and Johnson 2013, Yang, Wills et al. 2015)**. To find the optimal conditions to conduct our screen, and to understand how SCβ cells respond to stress, we performed an initial stress assay in which we exposed SCβ cells to various culture conditions. We differentiated and cultured *INS*-EGFP stem cell derived cells to day 35, dispersed the spheroids into single cells, treated them with a total of 48 conditions, stained them with Hoechst cell nuclei marker and propidium iodide (PI) cell death marker, and performed live cell imaging (Figure 4A,B; Supplemental Figure 7A-D, 8A). Conditions selected reflected possible challenges that SCβ cells are likely to encounter in the event of a transplant, including glucose concentrations (hyperglycemia or hypoglycemia), thapsigargin (Tg) as a stimulator of ER stress, and cytokines to model immune reaction. We also tracked EGFP as a readout for *INS* gene activity**(Chu, Modi et al. 2022)** over 96 hours of imaging (Figure 4B, 5A-C). Tg induced cell death in a dose dependent manner, and that cell death was mildly but not significantly reduced in ITS-containing conditions (Figure 4C,D). We found that cell death was dose dependent with the cytokine cocktail (Figure 4E,F). Cell death was reduced with the addition of ITS, with significant effects in lower cytokine cocktail concentrations, but not in higher concentrations of cytokine cocktail (Figure 4E,F). After filtering out dead or dying cells based on PI fluorescence, we found that in most conditions, treatment with Tg or cytokines reduced *INS* gene activity (Figure 5A,B). However, we found that *INS* gene activity was primarily controlled by glucose concentrations, with higher glucose concentrations increasing EGFP fluorescence (Figure 5C). Together our data show that SCβ cell death can be induced dose dependently by stressors, and that SCβ cell *INS* gene activity could be manipulated by glucose concentrations.

**Figure 4.**
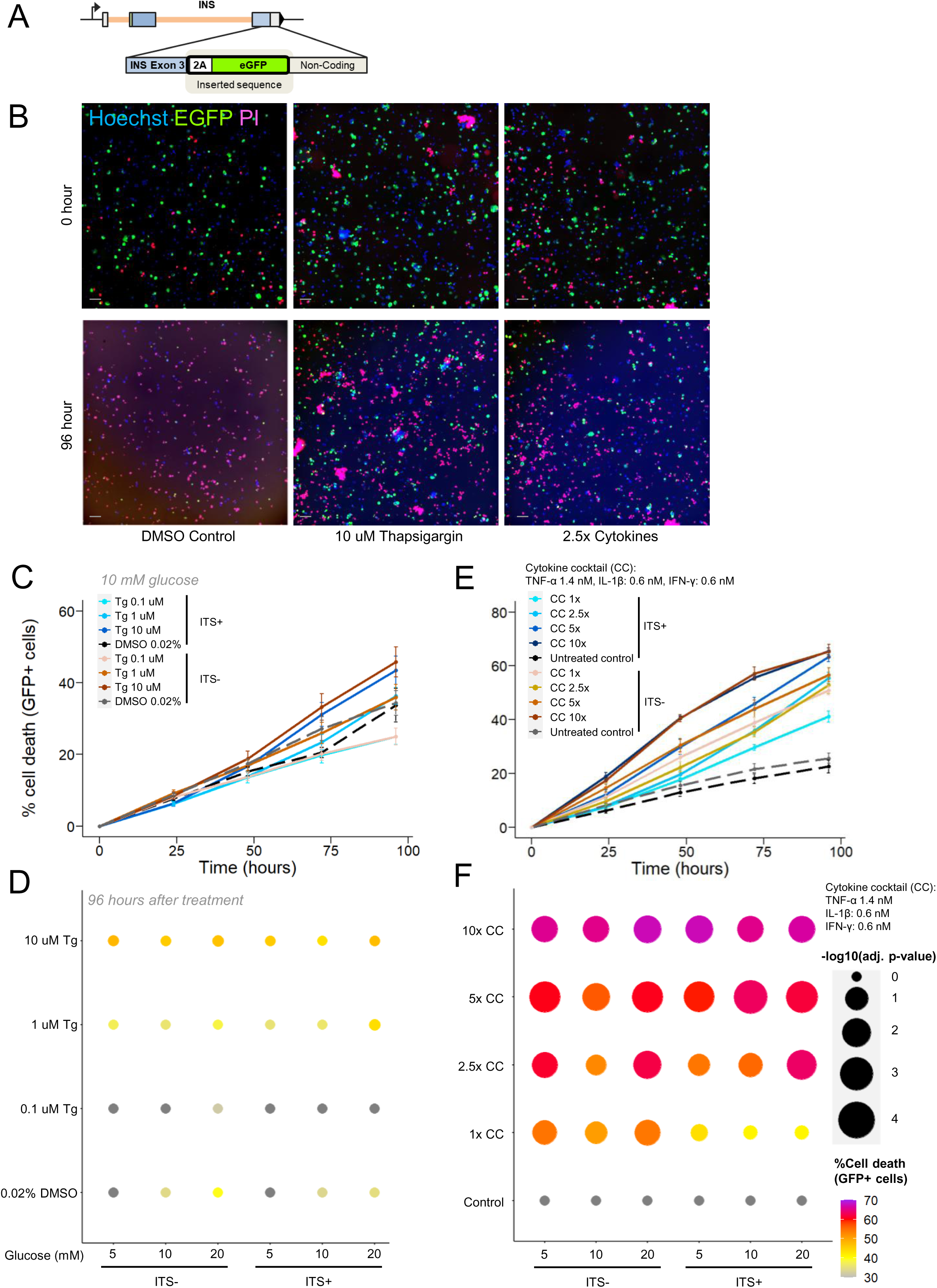
**Selecting stress conditions for a live imaging screen for SC**β **cells survival. (A)** Schematic depicting the *INS*-EGFP stem cell line. **(B)** Example images of the assay at 0 hours and 96 hours of imaging. Blue wavelength is Hoechst cell nuclei dye, red is the cell death marker propidium iodide, green is EGFP. Scale bar is 40 μm. **(C)** Survival curves depicting dose dependent cell death induced by various concentrations of thapsigargin at 10 mM glucose. Error bars are SEM. **(D)** Dot plot of EGFP positive cell death thapsigargin conditions after 96 hours of imaging. Size of dots denotes – log10 (adjusted *p* value ) compared to cytokine control. Color of the dots represent percentage of PI positive cells. **(E)** Survival curves depicting dose dependent cell death induced by various concentrations of cytokine cocktail at 10 mM glucose. Error bars are SEM. **(F)** Dot plot of EGFP positive cell death in cytokine cocktail conditions after 96 hours of imaging. Size of dots denotes – log10 (adjusted *p* value) compared to cytokine control. Color of the dots represent percentage of PI positive cells. n=3 differentiations. Two-way ANOVA.

**Figure 5.**
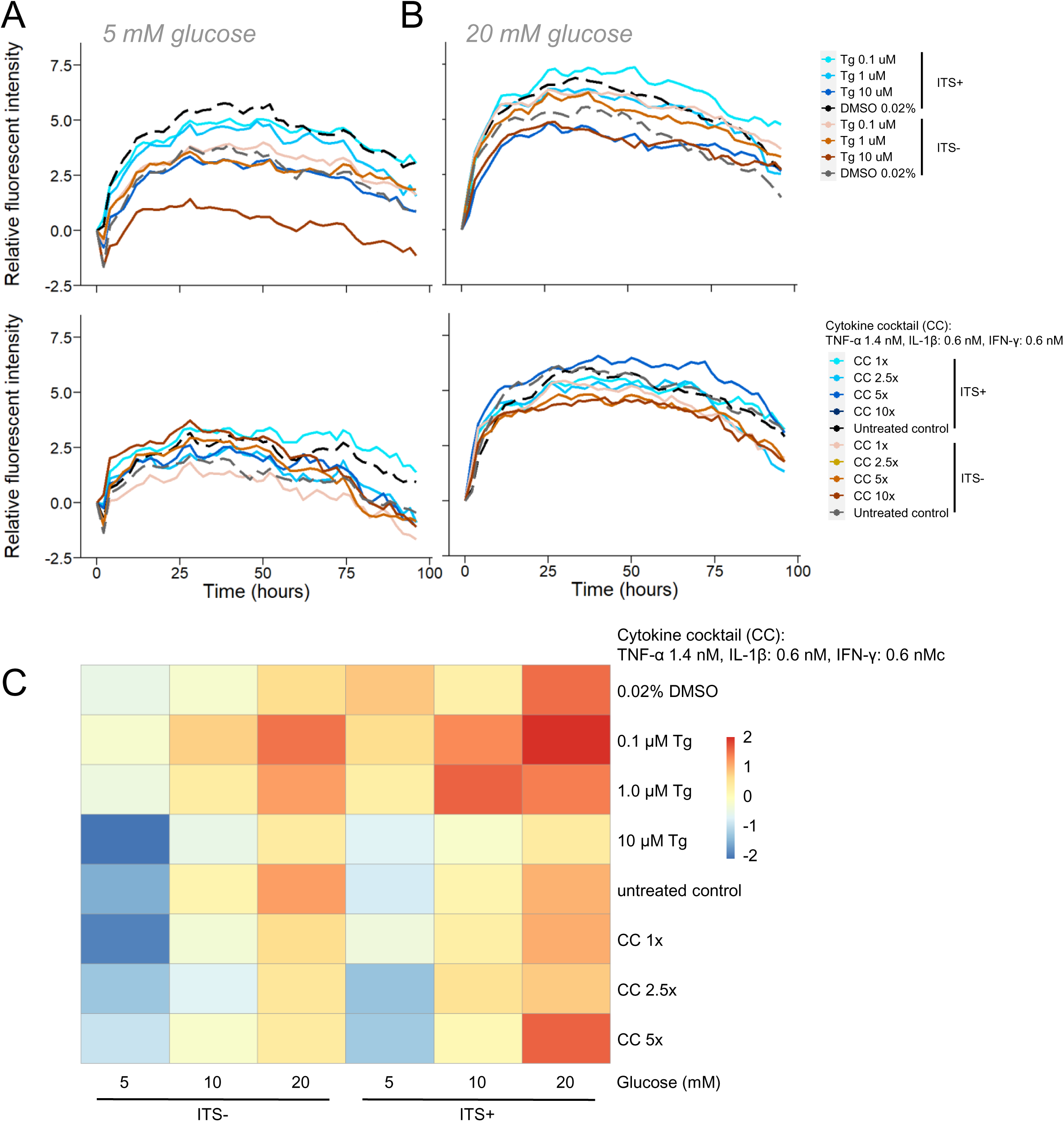
*INS* gene activity under stress conditions. **(A)** EGFP fluorescence over time in 5 mM glucose conditions in various doses of thapsigargin and cytokine cocktail. **(B)** EGFP fluorescence over time in 20 mM glucose conditions in various doses of thapsigargin and cytokine cocktail. **(C)** Heatmap of *INS* gene activity (AUC) over 96 hours of imaging across all conditions measured. n=3 differentiations.

### A high-throughput screen of 148 ligands for mediators of cell survival

We chose a condition that induced reliable but not excessive cell death (Figures 4F, 5C) for the screen of prioritized ligands (Figure 3F). Our rationale behind choosing 10 mM glucose, without ITS, 1.6 nM TNF-α, 0.7 nM IL-1β, was 0.7 nM IFN-γ was: 1) the cytokine conditions induced a consistently high level of cell death; 2) cytokine 1x conditions were the only cytokine conditions which displayed a clear response to the addition of ITS, suggesting that this condition is ‘rescuable’; 3) Out of the three cytokine 1x conditions, though the 20 mM glucose has the highest “rescue” effect, 20 mM glucose may be a bit too stimulatory, as shown with our EGFP fluorescence data. (Figure 4A-E, 5A-C). Overall, we reasoned that Cyto1x, 10 mM glucose without ITS was the best choice. For our choice of ligands, we decided to use the scarce ligands with abundant receptors list (Fig. 3E, F). This list represents ligands which are reduced in SC-islets compared to human islets but have abundant receptors which may be primed for activation. Similar to our stress condition pre-screen, we conducted 96-hour live cell imaging experiments on day 35 dispersed SCβ cells with 148 ligands from our prioritized list; we used EGFP as a readout for *INS* gene activity and PI as a cell death marker. Importantly, these experiments were repeated 5 independent times, providing robust results. Concentrations used were chosen from rigorous online research on prior studies which utilized these ligands. After 72 hours, several fibroblast growth factors (FGF5, FGF8F, FGF6, FGF4, FGF19) showed significant protective effects (Supplemental Figure 9A). On the other end of the spectrum, we found a number of interferons (IFNW1, IFNA14, IFNA16, IFNA17, IL28B) increased cell death compared to our cytokine control (Supplemental Figure 9A). After 96 hours of imaging, we observed continued protective effects from various FGFs, with FGF5, FGF19, and FGF8F being the standouts and having a final cell death percentage comparable to our untreated control (Figure 6A). We found a cluster of ligands with high cell death (over 80%) at 96 hours (Figure 6A). These data show that members of the FGF family of ligands had a strong protective effect against immune induced cell death, while the addition of interferons may exacerbate cytokine cocktail driven SCβ cell death.

**Figure 6.**
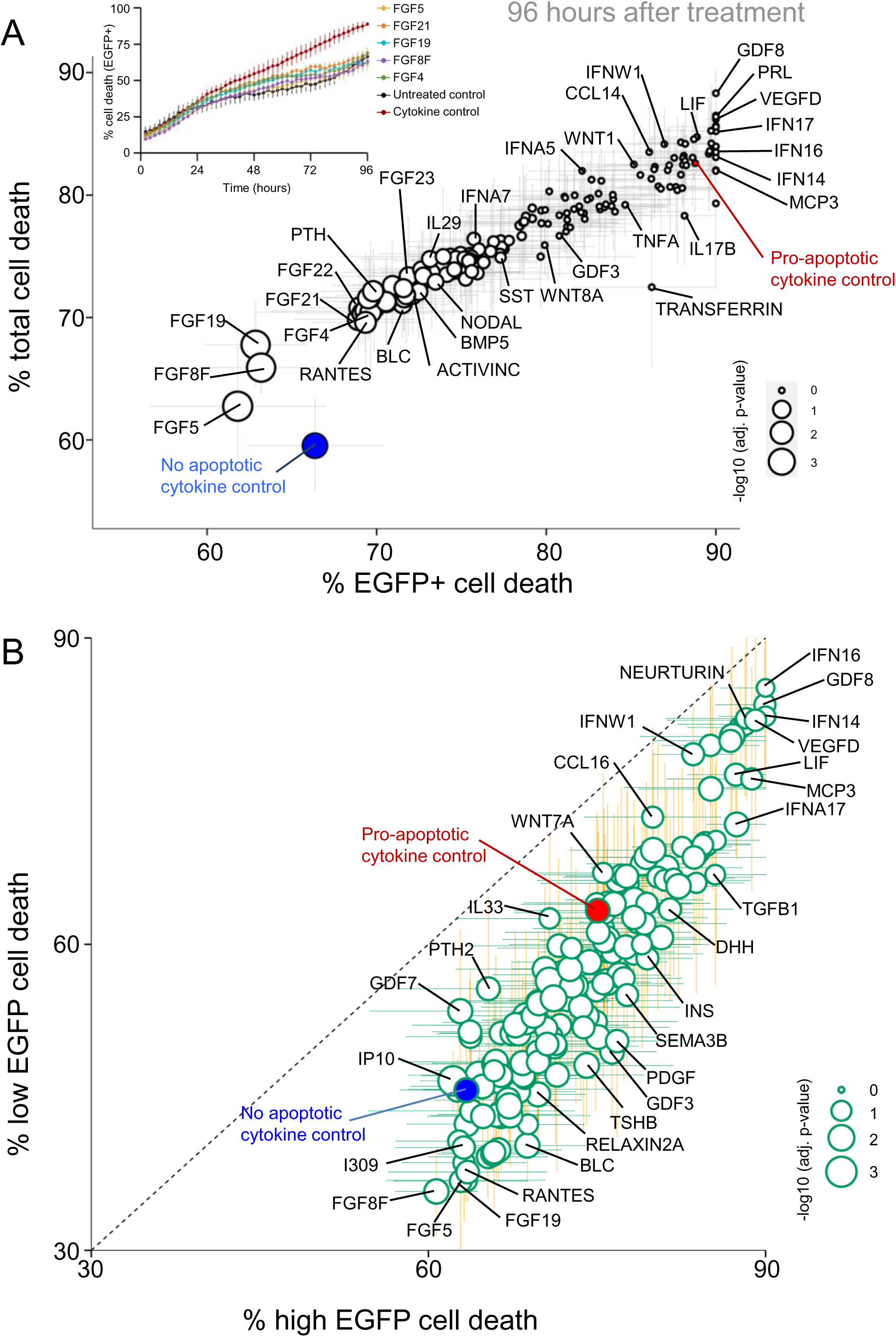
**Screening a prioritized list of 148 ligands for SC**β **cell survival in the context of pro-apoptotic cytokines. (A)** Total cell death in EGFP positive cells after 96 hours of treatment. Size of the dots denotes –log10 (adjusted *p* value) compared to pro-apoptotic cytokine control. Base condition for all treatments except untreated controls were 10 mM glucose, with 1x cytokine cocktail, without ITS. No apoptotic cytokine controls were 10 mM glucose, without ITS, without cytokine cocktail. One-way ANOVA. **(B)** Cell death of cells with low EGFP fluorescence against cells with high EGFP fluorescence after 72 hours of treatment. Error bars are SEM. Cells were tracked individually overtime. n=5 differentiations. Size of dots denotes –log10 (adjusted *p* value) when comparing high EGFP cell death to low EGFP cell death within each condition. n=5 differentiations. Paired t-test adjusted for multiple comparisons.

We next investigated *INS* gene activity in the context of our stress condition. We only included data from cells that stayed alive throughout the experiment. As expected, our untreated IBMX positive control had the highest levels of EGFP fluorescence, consistent with effects we have previously shown(Chu, Sabbineni et al. 2025) (Supplemental Figure 9B). At 72 hours, compared to the cytokine cocktail control, the strongest upregulators of *INS* gene activity were FGF4, FGF5, FGF6 and FGF8E, while the most significant downregulators were EPO, MCP3, and VEGFD. This trend suggested that members of the FGF family could reverse the detrimental effects of the cytokine cocktail on *INS* gene activity (Supplemental Figure 9B). Overall, these data suggest that members of the FGF family protect SCβ cells against cytokine-induced apoptosis and could rescue insulin production dysfunction under stress conditions. These effects were not associated with evidence of increased proliferation in our 96-hour imaging experiments (Supplemental Figure 10C).

We previously showed that mouse β cells with higher *Ins2* gene activity were significantly more vulnerable than cells with lower insulin production(Chu, Modi et al. 2022, Chu, Sabbineni et al. 2025). Therefore, we split the cells in our screen into two categories based on EGFP fluorescence and reanalyzed the data. Strikingly, we found that SCβ cells with higher EGFP were more vulnerable to the cytokine cocktail compared to cells expressing lower EGFP, consistent with the data from mouse primary β cells (Figure 6B). Interestingly, the top FGF hits at 96 hours (FGF5, FGF8F, and FGF19) had stronger protective effects on cells with low EGFP fluorescence even when compared to our untreated control (Figure 6B). This suggests that the FGFs have preferential protection on cells with lower *INS* gene activity.

### Dose-dependent rescue effects of top FGFs

We next selected the top FGF hits (FGF4, FGF5, FGF8F, FGF19, FGF21) and performed an independent dose-response experiment (0.01, 0.1, 1, 10, 100 nM) on two separate cell lines to confirm our findings and compare their protective potency (Figure 7A). The protective effects of all five FGFs were dose dependent in the *INS*-EGFP cell line, with a bit more variability in the HAF-iPSC cell line (Figure 7A ,B). Consistently across both lines, FGF4 had strong protective effects across all doses and FGF8F showed the strongest dose-dependency (Figure 7C-F). FGF19 and FGF21 did not show strong protective effects in the HAF-iPSC cell line (Figure 7B), although the lack of a INS-GFP reporter meant that these results were on mixed populations of cells and may not reflect specific effects on β cells. We also analyzed EGFP fluorescence in live SCβ cells and found that higher doses of FGF4, FGF5, and FGF8F had stronger rescue effects when compared with the cytokine control (Supplemental Figure 11A). Protein translation is altered under stress conditions in β cells. Thus, we measured translational capacity using O-propargyl-puromycin incorporation into nascent proteins. Although not statistically significant, FGF4 had a dose-dependent effect on protein translation in this assay (Supplemental 11B). These experiments provide strong rationale for future studies aimed at examining the mechanisms by which selected FGF ligands protect SCβ cells.

**Figure 7.**
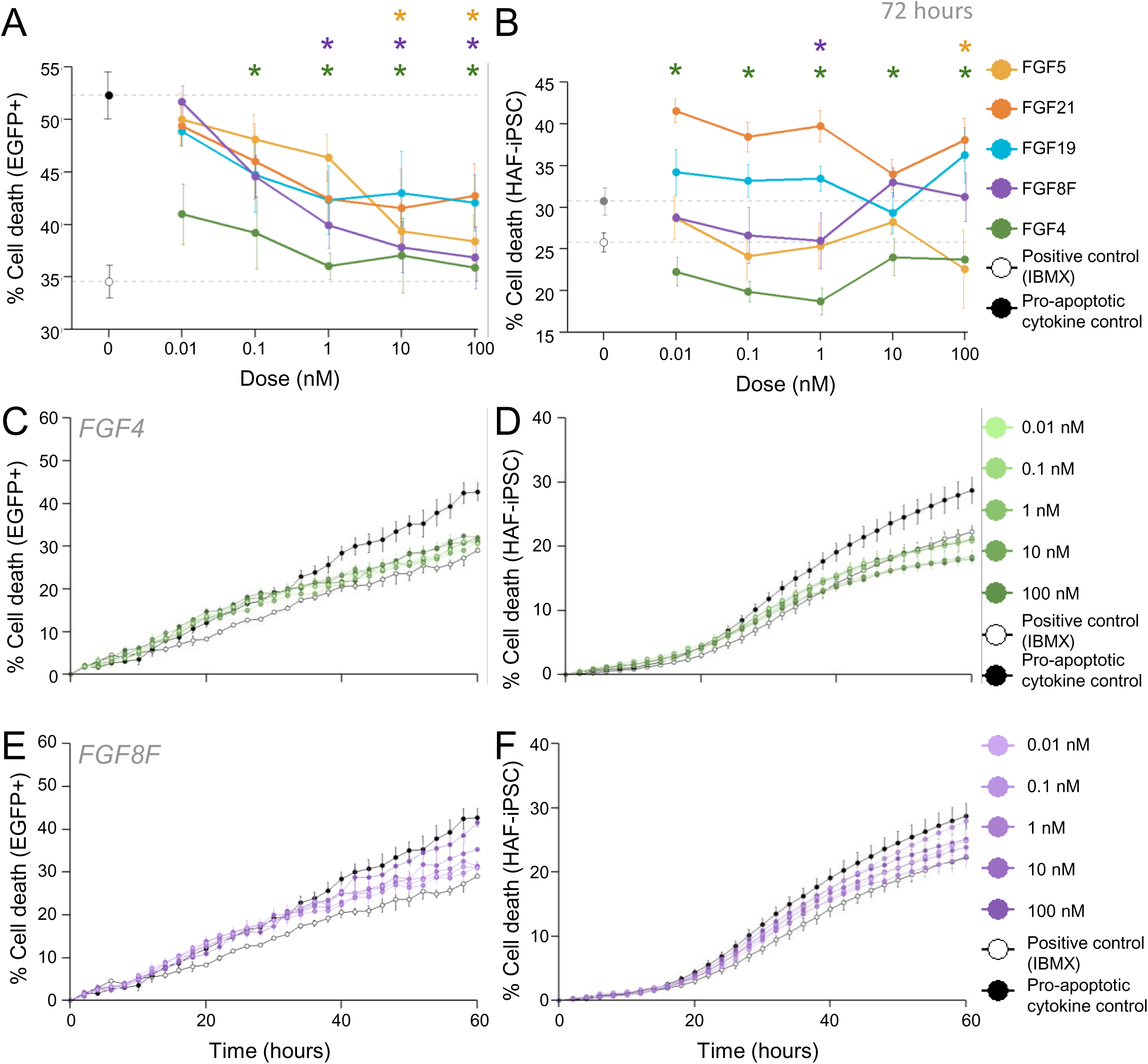
Dose-dependent protective effects of FGF ligands. **(A)** Percent EGFP positive cell death at a range of doses (0.01, 0.1, 1, 1, 10 nM) in the context of cytokine induction after 72 hours. n=3 differentiations. **(B)** Percent cell death in HAF-iPSC cells at a range of doses (0.01, 0.1, 1, 1, 10 nM) in the context of cytokine induction after 72 hours. n=5 differentiations. (C) Percent EGFP positive cell death against time in the context of doses of FGF4. n=3 differentiations. (D) Percent cell death in HAF-iPSC cells against time in the context of doses of FGF4. n=5 differentiations. **(E)** Percent EGFP positive cell death against time in the context of doses of FGF8F. n=3 differentiations. (F) Percent cell death in HAF-iPSC cells against time in the context of doses of FGF8F. n=5 differentiations. Base condition for all treatments except untreated controls were 10 mM glucose, with 1x cytokine cocktail, without ITS. Untreated controls were 10 mM glucose, without ITS, without cytokine cocktail. n=3 differentiations. Asterisks represent adjusted *p* value < 0.05. One-way ANOVA.

## Discussion

In this study, we systematically mapped intercellular signaling in human islets and SC-islets, built prioritized lists of ligands and receptors in SC-islets, and identified factors which protect SCβ cell under stress conditions. Our list of 422 protein ligands, 349 receptors, and 1552 interactions is the first reported analysis of intercellular communication in islets to combine proteomics, bulk RNA-seq, scRNA-seq, and novel ligand-receptor resources. Using this list and long-term high-content imaging, we identified multiple members of the FGF family which exert strong protective effects against immune-induced SCβ cell death and confirmed the efficacy of FGF4, FGF5, and FGF8F in an independent stem cell line. These findings provide a pathway to producing more resilient SCβ cells for type 1 diabetes therapy.

Intercellular signals play critical roles in adult islet function, and their dysregulation contributes to diabetes pathophysiology(Brun, Grijalva et al. 2015, Jin, Guerrero-Juarez et al. 2021). Multiple studies have also shown that unknown soluble ligands from the host promote rapid in vivo maturation of transplanted SCβ cells(Rezania, Bruin et al. 2014, Velazco-Cruz, Goedegebuure et al. 2020, Augsornworawat, Maxwell et al. 2021). Our results highlighted the contributions of α-cells, pericytes, duct cells, and endothelial cells to intra-islet signaling. While paracrine signals from α-cells are well studied(Rodriguez-Diaz, Molano et al. 2018, Huising 2020, Rodriguez-Diaz, Tamayo et al. 2020), paracrine signals from these other islet cell types have only recently been identified(Aamodt and Powers 2017, Saunders, Aamodt et al. 2021). Pericytes have been shown to promote β cell function and maturation by secreting BMP4(Houtz, Borden et al. 2016, Sasson, Rachi et al. 2016, Sakhneny, Rachi et al. 2018). Interestingly, we did not detect BMP4 in our human islet scRNA-seq and found BMP signaling occurred in the opposite direction; β cells sent signals to pericytes via BMPs. The data we presented provide direction for future studies to elucidate the roles of these non-endocrine cells in islet paracrine signaling. Bosi et al. used CellPhoneDB and single-cell transcriptomics to identify 9707 interactions in human islets from non-diabetic donors(Bosi, Marselli et al. 2022). We identified 6557 inferred interactions in human islets using a single-cell dataset containing 10x more cells. However, a direct comparison of the number of interactions between studies is unlikely to be meaningful due to inherent differences in the datasets and analysis tools between studies. Bosi et al. identified EGFR, FGFR1, and FGFR2 as top receptors involved in multiple interactions in human islets. Interestingly, while we did not analyze individual ligand-receptor pairs in human islets, we identified EGFR and FGFR1 as top abundant receptors with abundant ligands in SC-islets. These receptors have well-established roles in islet cell survival and development(Arany and Hill 2000, Hart, Baeza et al. 2000, Miettinen, Huotari et al. 2000). To create non-redundant prioritized ligand and receptor lists, we selected only the highest-ranked interaction for ligands and receptors with multiple interactions. In contrast, Bosi et al. used the number of interactions associated with each ligand or receptor as a key component of their ranking system. Our results may have differed if we had used a similar approach. We identified the MK pathway as the top pathway to intra-islet signaling in both human islets and SC-islets. We also found that MDK, the main MK pathway ligand, was the top upregulated ligand at the protein level in SC-islets compared to human islets. Interestingly, a recent study also identified MK as one of the top three contributing pathways in the developing fetal human pancreas(Sean dlO 2022). MDK is known to play a role in embryogenesis and is upregulated in various cancers(Filippou, Karagiannis et al. 2020), however its role is islet biology is unknown. Future studies could examine the roles of MDK in human islets and SC-islets.

We generated three prioritized lists of interactions in SC-islets to guide future studies. We found FGF21 was in three of the top five interactions between scarce ligands and abundant receptors in SC-islets, indicating FGF21 could be a “missing” ligand. FGF21 has been shown to improve β cell survival and function(Wente, Efanov et al. 2006), protect against type 2 diabetes in mice(Pan, Wang et al. 2019), and improve islet engraftment following islet transplantation in mice. Interestingly, a recent study showed that FGF21 from the liver is involved in β cell regeneration(Cui, Feng et al. 2023). Additional studies on the role of FGFs, including in primary islet cells, are warranted.

The effects of stress on primary β cells are well documented(Harding and Ron 2002, Papa 2012, Yang, Wills et al. 2015, Dludla, Mabhida et al. 2023), but less was known about the effects of ER stress and cytokine-induced stress on stem cell-derived insulin producing cells(Demine, Schiavo et al. 2020, Castro-Gutierrez, Alkanani et al. 2021, Dettmer, Niwolik et al. 2022, Leite, Pelayo et al. 2022). Here, we provide a comprehensive study with multiple doses of stress-inducing conditions on SCβ cells with temporal information. We show that SCβ cell death in response to thapsigargin and cytokine cocktails are dose-dependent. Prior studies have shown that treatment with cytokines on INS-1E and human islet cells induces differential gene expression after 24 hours and 48 hours of treatment respectively(Lopes, Kutlu et al. 2014). Interestingly, we found that at the 24-hour time-point, a major increase in cell death is observed which is reduced after 48 hours; this may coincide with the differential gene expression observed in the aforementioned study(Lopes, Kutlu et al. 2014). *INS* gene activity was reduced by stresses and influenced by glucose levels. We have previously shown how endogenous *Ins2* gene activity can be influenced by glucose and serum levels(Chu, Modi et al. 2022).

From our prioritized ligand list, we identified several members of the FGF family of ligands as protective against cytokine-induced apoptosis. Our ligand screen and dose response experiments revealed that FGF19, FGF21, FGF5, and FGF8F had protective effects. FGF4 had strong protective effects at all doses measured. These five ligands were from three FGF subfamilies, the FGF4 subfamily (FGF4, FGF5), the FGF8 subfamily (FGF8F), and the ‘endocrine’ FGF19 subfamily (FGF19, FGF21). FGF6, the last member of the FGF4 subfamily, also had strong protective effects but they did not persist to 96 hours. Members of the FGF19 subfamily are considered endocrine FGFs and are known to play roles in mature islets in mice, including protection against cytokine-induced stress(Pan, Wang et al. 2019, Edirisinghe, Ternier et al. 2024). FGF21 has been shown to improve INS-1E and rat islet function and survival via signalling of the extracellular signal–regulated kinase 1/2(ERK 1/2) and Akt pathways(Wente, Efanov et al. 2006). While not directly tested on insulin-producing cells, a study identified a positive relationship between FGF19 serum levels and β cell function in humans(Tang, Su et al. 2019). Interestingly, of our four FGF8 isoforms (FGF8A, FGF8B, FGF8E, FGF8F) only FGF8E and FGF8F had protective effects. Due to FGF8E and FGF8F isoforms having only recently appeared in placental mammals, it has been suggested that they may possess distinct functions compared to FGF8A and FGF8B(Sunmonu, Li et al. 2011). The primary difference between the earlier and later isoforms is the 1C exon, a region where mutations have been known to result in idiopathic hypogonadotropic hypogonadism, a disease associated with deficient gonadotropin releasing hormone (GnRH)(Falardeau, Chung et al. 2008). More studies are needed to investigate the role of these FGF8 isoforms, possibly with in silico studies using AlphaFold to identify key binding regions in the 1C exon(Abramson, Adler et al. 2024). Members of the FGF4 family have been shown to play roles in embryonic development and tissue repair, possibly explain their strong protective effects when compared to the other FGFs(Kosaka, Sakamoto et al. 2009). FGF4, in particular, has been shown to play roles in stem cell differentiation and proliferation(Johannesson, Stahlberg et al. 2009, Kosaka, Sakamoto et al. 2009). Validation experiments on the HAF-iPSC cell line revealed that members of the FGF19 family (FGF19, FGF21) did not have significant protective effects, suggesting that some FGF effects are cell line-specific. Our study is the first to identify members of the FGF4 family as having protective effects in the context of cytokine-induced stress. However, as would be expected of the less mature SCβ cells, there are discrepancies when comparing our findings with those in mature primary β cells and cell lines. Overall, our study is the first to show that some of the effects in FGFs can extend to SCβ cells, and we also identify novel members of the FGF subfamilies which may modulate SCβ cell survival.

In summary, we integrated multiple online resources and databases and investigated the cell-cell interactions in primary human islets and SC islets. We identified and ranked the top interactions in human primary islets and compared these interactions to those in SC islets. We then generated a prioritized list of ligands based on the discrepancies between human islets and SC islets and conducted a targeted image-based screen to identify factors that can protect SCβ cells from stresses associated with the immune attack that might occur following transplantation or during type 1 diabetes. We identified multiple members of the FGF ligand family that exhibited reproducible protection in the context of cytokine-induced apoptosis. These results reveal possible pathways to target in SCβ cells to alleviate the stress of immune cell-mediated death in cell replacement therapy. This work is an important step towards understanding the differences between human β cells and stem cell derived insulin producing cells, and advancement towards a cell therapy for diabetes.

## Resource availability

### Lead contact

Requests for further information and resources should be directed to and will be fulfilled by the lead contact, James D. Johnson.

### Materials availability

This study did not generate new unique reagents.

### Data and code availability

scRNAseq data have been deposited at Gene expression omnibus as GEO:GSM9888508 and GEO:GSM9888509 and are publicly available as of the date of publication.

R coding have been deposited at Github as Github: 932224975 and are publicly available as of the date of publication.

## Acknowledgments

We thank many colleagues for helpful discussions. This manuscript used data acquired from the Human Pancreas Analysis Program (HPAP-RRID:SCR_016202) Database (https://hpap.pmacs.upenn.edu), a Human Islet Research Network (RRID:SCR_014393) consortium (UC4-DK-112217, U01-DK-123594, UC4-DK-112232, and U01-DK-123716).

## Author Contributions

CMJC designed studies, performed experiments, analyzed/interpreted data, wrote manuscript. SK designed studies, analyzed/interpreted data, helped write manuscript. LTH designed and performed experiments. CLB, MJB, SM designed and performed experiments, analyzed/interpreted data. SYC performed experiments and helped write the manuscript. MEO designed studies, analyzed and interpreted data. WWW, FCL, JDJ conceived the project, designed studies, interpreted data, edited the manuscript, and obtained funding. FCL and JDJ are the guarantors of this work.

## STAR Methods

### Experimental model and study participant details

#### Cell lines

A human embryonic stem cell line (INS-2A-EGFP) that contains a knock-in add-on of EGFP downstream was derived from WA01 (hPSCReg #WAe001-A) (Novakovsky, Sasaki et al. 2023, Kolic, Sun et al. 2024, Mar, Filatov et al. 2025). This INS-2A-EGFP clone source is from the Lynn lab, and has not been registered. Another cell line, the HAF-iPSC cell line (hAF12, official name CHUSJ-012), was obtained from The Human Health Therapeutics Research Centre, National Research Council of Canada, Ottawa, ON (Hwang, Khalil et al. 2026). This cell line has also not been registered. The sex of both cell lines are XY. Cells were grown in mTESR Plus on Geltrex, and split at 90% confluence with ReLeSR or Accutase at ratio of 1:10-1:15. Passage numbers for INS-2A-EGFP were between 40-55, while passage numbers for the HAF-iPSC line were between 20-30. Authentication of cell lines were done by karyotyping with a 28 probe ddPCR at StemGenomics and found free of common abnormalities, and the cells were tested monthly for mycoplasma contamination by qPCR and found to be free of contamination.

*Ethics statement*

The WA01 hESC line was generously provided by WiCell (Madison, WI, USA); HAF-012 iPSC line was provided by the NRC. Experiments were approved by the University of British Columbia (UBC) / Children’s and Women’s Health Centre of British Columbia Research Ethics Board (hESCs: H09-00676) and the CIHR Stem Cell Oversight Committee.

### Method details

#### Differentiation of stem cell derived spheroids

After the initial 21 days of differentiation, SCβ cells were cultured in CMRL with 5.5 mM glucose (Thermo Fisher Scientific) containing 1% fatty acid-free BSA (Proliant), 1:100 Glutamax, 1:100 NEAA, 1 mM pyruvate, 10 mM HEPES, 1:100 ITS (Thermo Fisher Scientific), 10 μg/ml of heparin sulfate, 1 mM N-acetyl cysteine, 10 μM zinc sulfate, 1.75 μL 2-mercaptoethanol, 2 μM T3 and 0.155 mM ascorbic acid (Sigma Aldrich) until Day 27. Between Day 27-35, aggregates were grown in CMRL with 5.5 mM glucose containing 1% fatty acid-free BSA, 1:100 Glutamax, 1:100 NEAA, 1 mM Pyruvate, 10 mM HEPES, 1:100 ITS, 10 μg/ml heparin sulfate, 1 mM N-acetyl cysteine, 10 μM zinc sulfate, 1.75 μL 2-mercaptoethanol, 10 nM T3, 1:2000 Trace elements A (Cellgro), 1:2000 trace elements B (Cellgro), 1:2000 Lipid Concentrate (Thermo Fisher Scientific) and 0.5 µM ZM447439 (Selleckchem)(Balboa, Barsby et al. 2022, Kolic, Sun et al. 2024).

For dispersion of stem cell derived spheroids into individual SCβ cells, spheroids were hand-picked and washed 2 times with PBS. Spheroids were then immersed for 8 minutes in Accutase (STEMCELL Technologies, Vancouver, Canada), with gentle flicking every 2 minutes. 4x the amount of PBS was then added to halt Accutase activity, and the cells were spun down at 200g, 5 minutes. Liquid was aspirated, and Day 27-35 media with 10µM ROCK inhibitor Y-27632 (STEMCELL technologies, Vancouver, Canada) was added.

#### Curated global lists of ligands, receptors, and interacting pairs

Supplemental Figure 1 provides an overview of this study’s methods, data inputs, and data outputs. Ligand, receptor, and interaction data were downloaded from the OmniPath web service using OmniPathR version 3.3.20 (https://github.com/saezlab/OmnipathR)(Turei, Korcsmaros et al. 2016, Turei, Valdeolivas et al. 2021). Ligand and receptor lists were obtained using the import_omnipath_intercell function. Protein complexes and non-secreted ligands were excluded. Ligand-receptor interactions were obtained using the import_omnipath_interactions function. Duplicate genes corresponding to the same HGNC symbol were merged. Ligands, receptors, and interactions without literature support were excluded.

Ligand and receptor lists were filtered in two steps. The OmniPath consensus score, which is the number of resources in the OmniPath database supporting the classification of an entity into a category, was used to filter out ligands and receptors that were only present in a few resources. The consensus score thresholds were chosen based on the distribution of consensus scores and the point at which the majority of the genes seemed to be true ligands and receptors based on our knowledge. Next, the remaining ligands and receptors were manually reviewed to determine whether they were appropriate for inclusion. Manual review was done by examining the OmniPath category, Entrez gene summary, OmniPath interaction partners, and gene ontology annotations for each gene. As required, literature references were also reviewed. Ligand and receptor pathway annotations were obtained from CellChat via OmniPath.

#### Bulk mRNA expression data

Three bulk RNA-seq datasets were used in this study. We were provided processed data outputs from Kallisto for the Lund 2014 human islet dataset (n=89), which contained islet donors of varying diabetes status (GEO: GSE50244)(Fadista, Vikman et al. 2014). Transcripts per million (TPM) was used to quantity gene expression to allow for accurate between-gene expression comparisons(Zhao, Ye et al. 2020). Gene level TPM values were calculated by taking the sum of all transcripts corresponding to the same Ensembl gene. Some HGNC symbols corresponded to multiple Ensembl genes. The highest TPM value corresponding to the HGNC symbol was used in these cases. We were provided TPM values for the MacDonald-Gloyn 2023 human islet dataset, which included 82 non-diabetic and 8 type 2 diabetic donors (n=90)(Kolic, Sun et al. 2024). For both these datasets, the mean TPM across all donors was used. To determine islet gene expression specificity, islet gene expression was compared to the expression across 54 human tissues from the Genotype-Tissue Expression (GTEx) Project(Consortium 2013, Mele, Ferreira et al. 2015). Median gene-level TPM by tissue from GTEx Analysis V8 was downloaded via the GTEx portal. The highest TPM value corresponding to each HGNC symbol was used.

*Proteomic data*

We used protein abundance data generated using liquid chromatography with tandem mass spectrometry from 133 human islet donors (available on HumanIslets.com(Ewald, Lu et al. 2024)) and SC-islets (n=18)(Kolic, Sun et al. 2024). 90 human islet donors were donor-matched to the MacDonald-Gloyn 2023 bulk RNA-seq dataset(Kolic, Sun et al. 2024). The SC-islets had been generated from an *INS*^EGFP/WT^ reporter human embryonic stem cell line using a stage 7 protocol and were a combination of S7 unsorted SC-islets (n=12) and S7 FACS-sorted SC-islets (n=6). The mean protein abundance was used for each group. The abundances were normalized to protein amino acid lengths obtained from UniProt to allow for accurate between-protein abundance comparisons. For proteins with multiple amino acid lengths, the mean length was used.

We used differential protein abundance analysis between the non-diabetic human islets (n=118) and pooled SC-islets (n=18). The data from the S7 unsorted SC-islets and S7 sorted SC-islets were pooled into a single SC-islet group. A protein was considered significantly differentially abundant if the absolute log2-fold change was >1 and the adjusted *p* value was <0.05. Only the 90 human islet donors with matching proteomics and transcriptomics data were used to determine the correlation between gene expression and protein abundance. The proteomics was compared to the MacDonald-Gloyn 2023 and Lund 2014 RNA-seq datasets. The linear correlation between protein and RNA was calculated using Pearson’s correlation coefficient.

#### Single-cell mRNA expression data

We used a published human islet scRNA-seq dataset generated from three cadaveric donors (n=32,486 cells, GEO:GSE196715)(Yoon, Sasaki et al. 2023). We also generated scRNA-seq datasets from SC-islets (n=6875 cells,GEO:GSM9888508) and FACS-sorted INS-GFP enriched, reaggregated SC-islets (n=5355 cells, GEO:GSM9888509). After INS-2A-EGFP knock-in, add-on hESCs were differentiated to SC-islet clusters (Stage 6 Day 21)(Novakovsky, Sasaki et al. 2023, Mar, Filatov et al. 2025), clusters were then dissociated, enriched for GFP-expressing cells using fluorescence-activated cell sorting (FACS), and reaggregated as previously described(Mar, Filatov et al. 2025). After another four days of culture (Stage 7 Day 25), unsorted and sorted clusters were washed with PBS dissociated using Accumax (STEMCELL Technologies, Vancouver, Canada). Cell suspensions were resuspended in PBS containing 10 μM Y-27632 (STEMCELL Technologies, Vancouver, Canada) and filtered to remove cell debris using a 35 µm nylon mesh (VWR, Radnor, United States). Library generation was performed according to manufacturer’s instructions for the Chromium Next GEM Single Cell 3’ Reagent Kit v3.1 (10x Genomics, Pleasanton, United States) as described(Yoon, Gamu et al. 2025). The data were processed using 10x Genomics Cell Ranger v6.1.1 and analyzed using Seurat version 4.1.0(Stuart, Butler et al. 2019, Hao, Hao et al. 2021, Hao, Hao et al. 2021). Differentially expressed genes with adjusted *p* values equal to zero were assigned adjusted *p* values equivalent to the lowest non-zero values. A gene was considered as significantly differentially expressed if the absolute log2-fold change was >1 and the adjusted *p* value was <0.05.

#### CellChat Intra-islet signaling analysis

Intra-islet signaling in human islets and S7 sorted SC-islets were analyzed using CellChat version 1.1.3 (https://github.com/sqjin/CellChat)(Jin, Guerrero-Juarez et al. 2021). The RNA assays from Seurat were used as inputs for CellChat. The “secreted signaling” subset of the default CellChat interaction database was used. Gene expression was projected onto the protein-protein interaction network to compensate for the potential dropout effects of signaling genes.

#### Ligand and receptor ranking in SC-islets

Three additional publicly available datasets were used in rankings. Bulk RNA-seq of S6 SC-islets (n=1) and human islets (n=1) from Alvarez-Dominquez et al was obtained from GEO (GEO: GSE139816)(Alvarez-Dominguez, Donaghey et al. 2020). A gene was considered differentially expressed in this bulk RNA-seq dataset if the absolute log_2_-fold change was >1. Expression of S6 SCβ cells and differential expression of S6 SCβ cells compared to human β cells from Veres et al., 2019(Veres, Faust et al. 2019)(GEO: GSM3141954). A gene was considered differentially expressed in this scRNA-seq dataset if the absolute log_2_-fold change was >1. Differential expression of S7 SCβ cells was compared to human β cells from Balboa et al., 2022 (n=3 differentiations, GEO: GSE167880)(Balboa, Barsby et al. 2022). A gene was considered differentially expressed in this scRNA-seq dataset if the absolute log_2_-fold change was >0.5 and the adjusted *p* value was <0.05.

Ligands and receptors were ranked in SC-islets based on gene expression or protein abundance, their relative expression or abundance relative to primary human islets (upregulation or downregulation), across 3 independent datasets. To increase the number of ranked gene product, the absolute log_2_-fold change threshold to be considered upregulated or downregulated was decreased to >0 for the proteomics and >0.5 for the Lynn scRNA-seq. Ligands and receptors were ranked on a percent scale for each parameter, with the top rank assigned a value of one and the bottom rank assigned a value of zero. Top abundant ligands and receptors were defined by high expression and upregulation ranks. Top scarce ligands and receptors were defined by low expression ranks and high downregulation ranks. The overall abundant rank and scarce rank for each ligand and receptor was calculated as the mean rank across the relevant parameters. Overall, the rankings were 50% based on expression in SC-islets and 50% based on differential expression compared to human islets.

Ligand and receptor ranks were combined to create three prioritized lists of interactions in SC-islets: abundant receptors with abundant ligands, scarce receptors with abundant ligands, and scarce ligands with abundant receptors. Interactions were ranked by calculating the product of the ligand and receptor ranks for each interacting ligand-receptor pair, as done previously(Efremova, Vento-Tormo et al. 2020). For receptors and ligands with multiple interactions, their highest ranked interaction partner was used for their final rank in the prioritized lists.

#### Live cell imaging and analysis

To image differential cell death in SCβ cells, day 35 *INS*-EGFP(Kolic, Sun et al. 2024) and HAF-iPSC stem cell derived spheroids were dissociated as above before being seeded onto Cultrex-treated (R&D systems, Minneapolis, United States) 384 well imaging plate (PerkinElmer, Waltham, United States) at 8000 cells per well. The next day, cells were washed 1 time with PBS before staining with Hoechst 33342 (0.05 μg/mL) and propidium iodide (0.5 μg/mL) for 2 hours. Treatments and ligands were then applied and live cell imaging carried out using an ImageXpress^Micro^ Confocal environmentally controlled, imaging system (Molecular Devices, San Jose, United States), which employed a Aura II light engine. Ligands and concentrations are listed in Table S1. Images were acquired with a 10x air objective (NA 0.3), at 2-hour intervals up to 96 hours, with cells being exposed to 359 nm light for 110 ms, 491 nm light for 15 ms, 561 nm for 75 ms. Analysis of live cell imaging experiments on *Ins2*^GFP^ cells was done using MetaXpress analysis software and custom R scripts. Tracking of GFP fluorescence in individual cells was performed using custom MetaXpress scripts and the *INS*(EGFP)^HIGH^ and *INS*(EGFP)^LOW^ subpopulations were identified using model-based clustering as previously described(Chu, Modi et al. 2022).

The matrix of conditions used in the initial stress assay included: 1) three doses of glucose (5, 10, 20 mM) to simulate hyperglycemia; 2) three doses of thapsigargin (Tg) to induce ER stress (0.1, 1, 10 μM); and 3) three doses of a cytokine cocktail comprising of tumour necrosis factor α (TNF-α), interleukin 1 β (IL-1β), and interferon γ (IFN-γ) (1.6 nM TNF-α, 0.7 nM IL-1β, 0.7 nM IFN-γ at 1x, 2.5x, 5x, 10x concentrations) to trigger immune factor induced stress; and 4) with and without the commercial insulin-transferrin-selenium (ITS) supplement. ITS is a key reagent used for SCβ cell culture and contains high concentrations of insulin, which is known to have protective effects on human β cells(Johnson, Bernal-Mizrachi et al. 2006). Proteins, their respective concentrations, and vendors used in the ligand screen are included in Table S1.

#### Protein synthesis assay

After 24 h of treatment, protein translation rates were measured using the Click-iT™ Plus O-propargyl-puromycin (OPP) Alexa Fluor™ 594 Protein Synthesis Assay Kit (Thermofisher, Cat. #C10457). Specifically, cells were incubated in growth media containing a 1:1000 dilution of reagent A for 30 min at 37°C, then washed in PBS, fixed in 3.7% formaldehyde for 15 min at room temperature, permeabilized with 0.5% Triton X-100 for 15 min at room temperature, and washed twice with PBS. Plates were stored at 4°C until the OPP assay was performed. The OPP assay was performed according to manufacturer instructions, except that volumes were scaled down for a 384-well plate, such that each well was incubated in 25 µL of reaction mix for 30 min in the dark at room temperature, washed in 25 µL of rinse buffer, then incubated in 25 µL of NuclearMask reagent (1:2000 in PBS) in the dark at room temperature. Finally, cells were washed four times with 50 µL of PBS and imaged at 10× with an ImageXpress Micro high-content imager using the DAPI (20 ms), Texas Red (120 ms), and FITC (180 ms) filters. Images were analysed with MetaXpress (Molecular Devices) to quantify the integrated staining intensity of OPP-Alexa Fluor 594 and EGFP fluorescence in cells identified by NuclearMask Blue Stain.

### Quantification and statistical analysis

All data analysis in used in generation of the prioritized list was performed with R version 4.1.2 in R Studio on a local computer. Statistics and data representation for live cell imaging experiments utilized custom R scripts. Student’s t-test, one-way ANOVAs, and two-way ANOVAs were used for parametric data analysis as indicated in Figure legends. For all statistical analyses, differences were considered significant if the p value was less than 0.05. Error bars were presented as ± standard error of the mean (SEM). Data practices conformed to FAIR and CARE principles

## Table S1

**List of ligands used for screen.** Includes ligand names, vendors, concentration used, and PMIDs of papers used to determine ligand concentrations for screen.

