## Supplemental figures and tables for "A prioritized medium-throughput screen identifies fibroblast growth factors as protective for human stem cell derived insulin secreting beta cells"

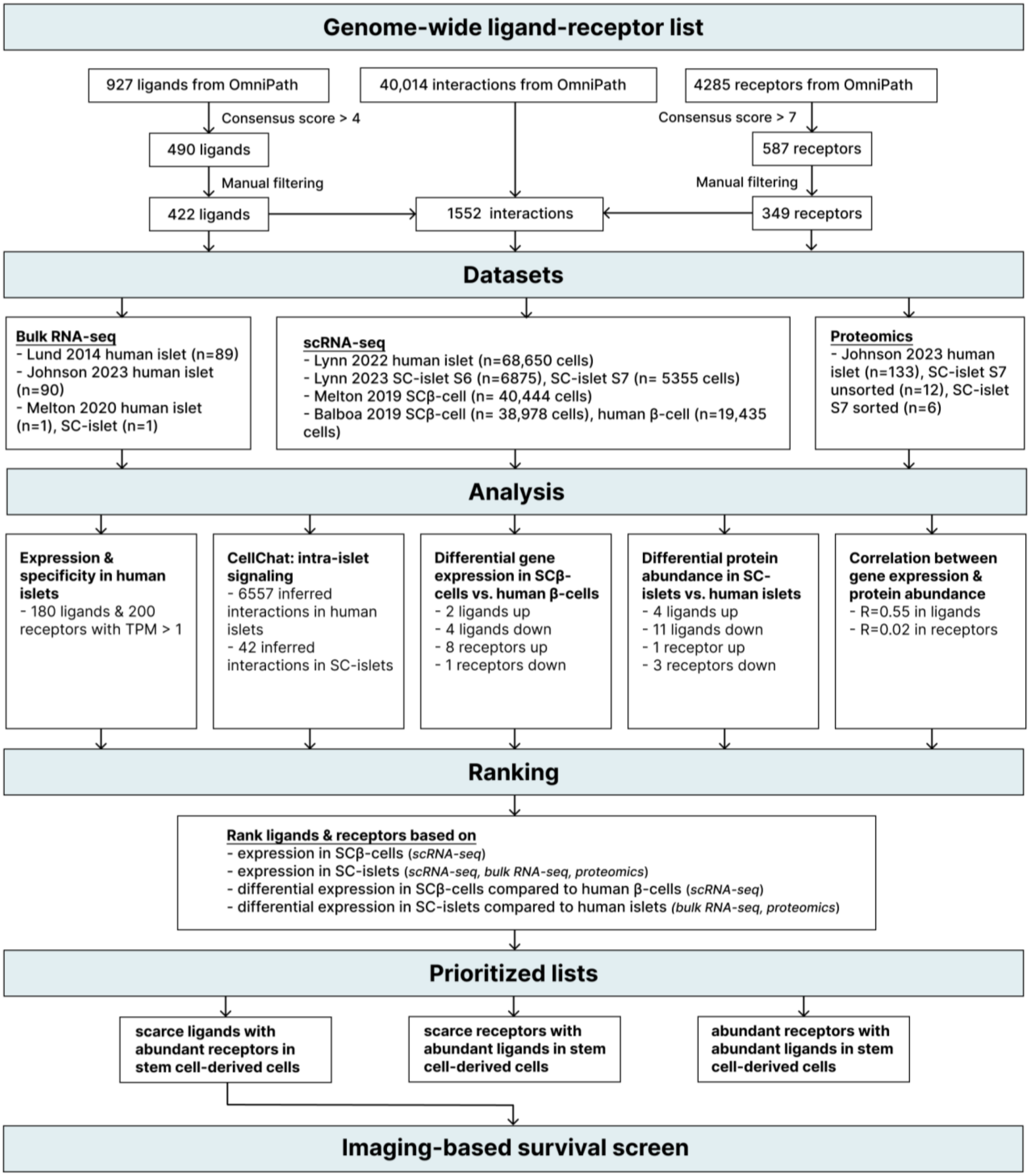


**Supplemental Figure 1. Process for production of the ligand list**. (A) Flowchart overview of methods, data inputs, and data outputs in this ligand list used in the screen.

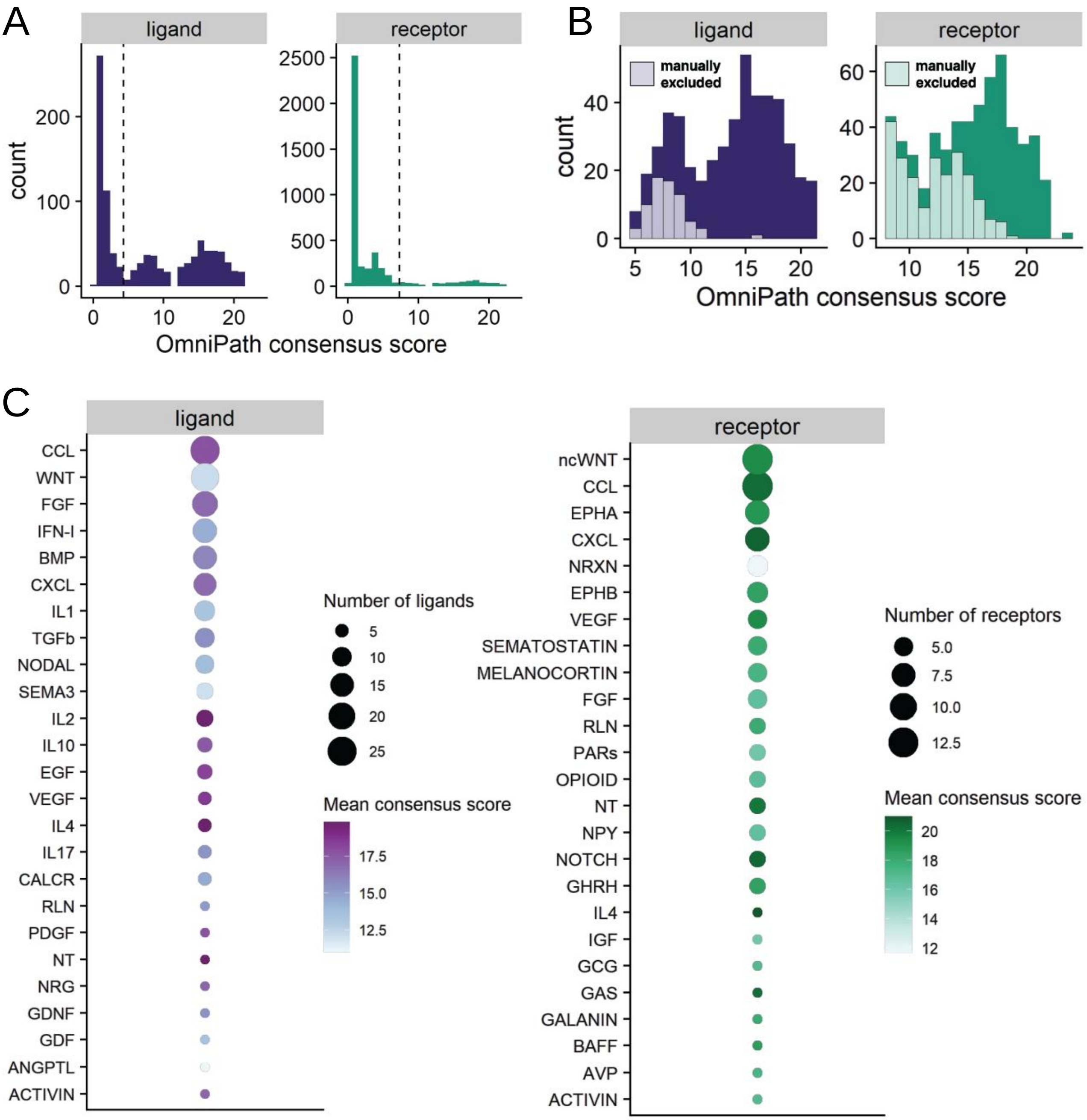


**Supplemental Figure 2. Semi-manual curation of 422 ligands and 349 receptors.**
**(A)** Distribution of OmniPath consensus scores for initial lists of 927 ligands and 4285 receptors obtained from OmniPath. Dashed vertical lines indicate the manually selected thresholds for inclusion. **(B)** Distribution of OmniPath consensus scores for manually excluded and included ligands and receptors. **(C)** Top 25 signaling pathways with the most ligands (left) and receptors (right) in the final lists. Dot size indicates the number of ligands or receptors corresponding to the pathway. The color indicates the mean consensus score across all ligands or receptors in the pathway.

A

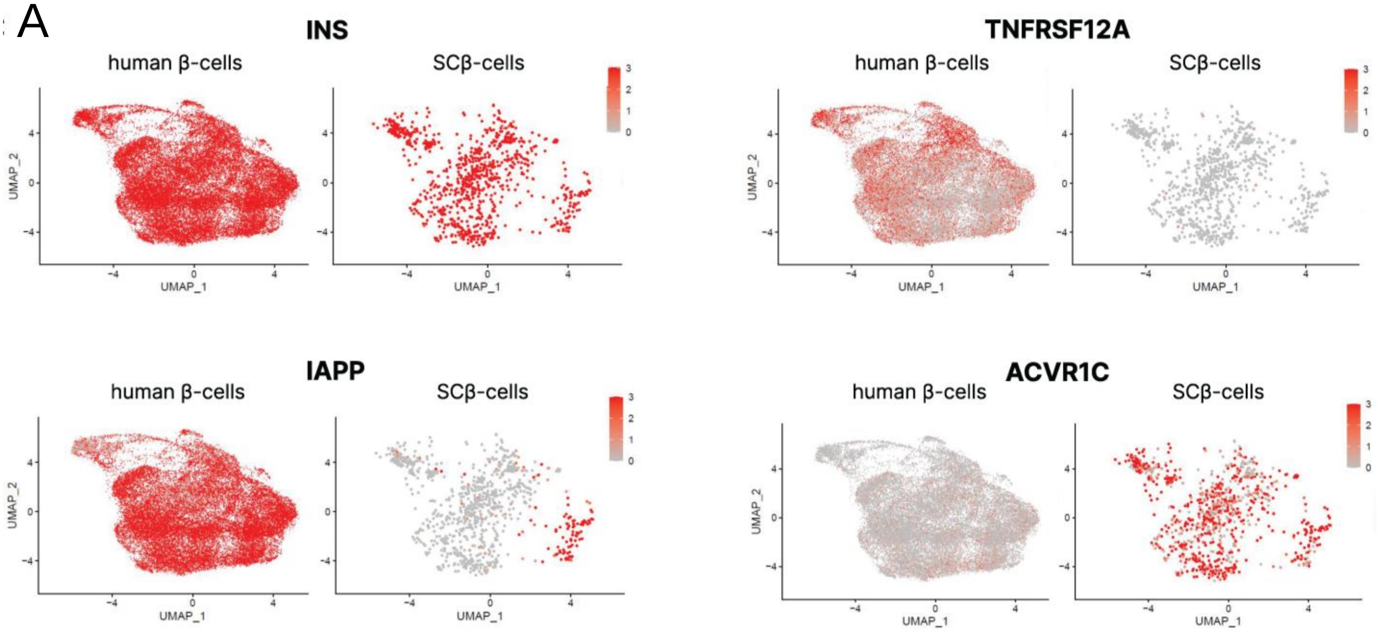


**Supplemental Figure 3. Example UMAPs of top genes**. **(A)** Single-cell gene expression of the top two differentially expressed ligands (INS & IAPP) and receptors (TNFRSF12A & ACVR1C) between SCβ-cells and human β-cells.

A

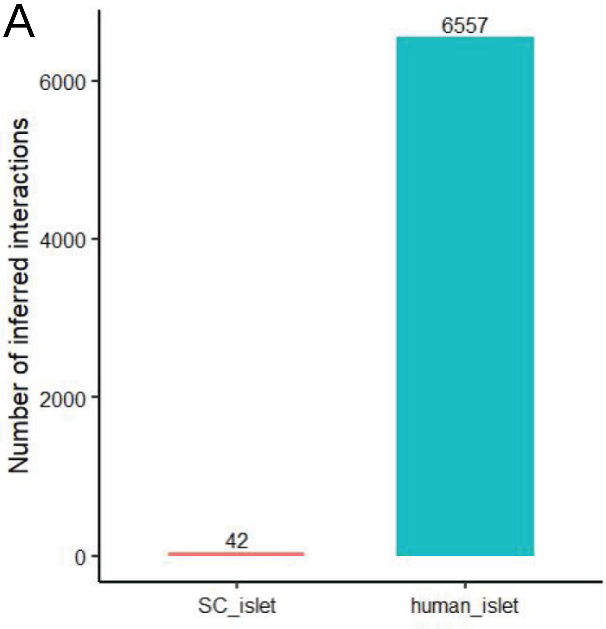


**Supplemental Fig 4. Inferred interactions. (A)** Total number of inferred interactions by CellChat in S7 sorted SC-islets and human islets.

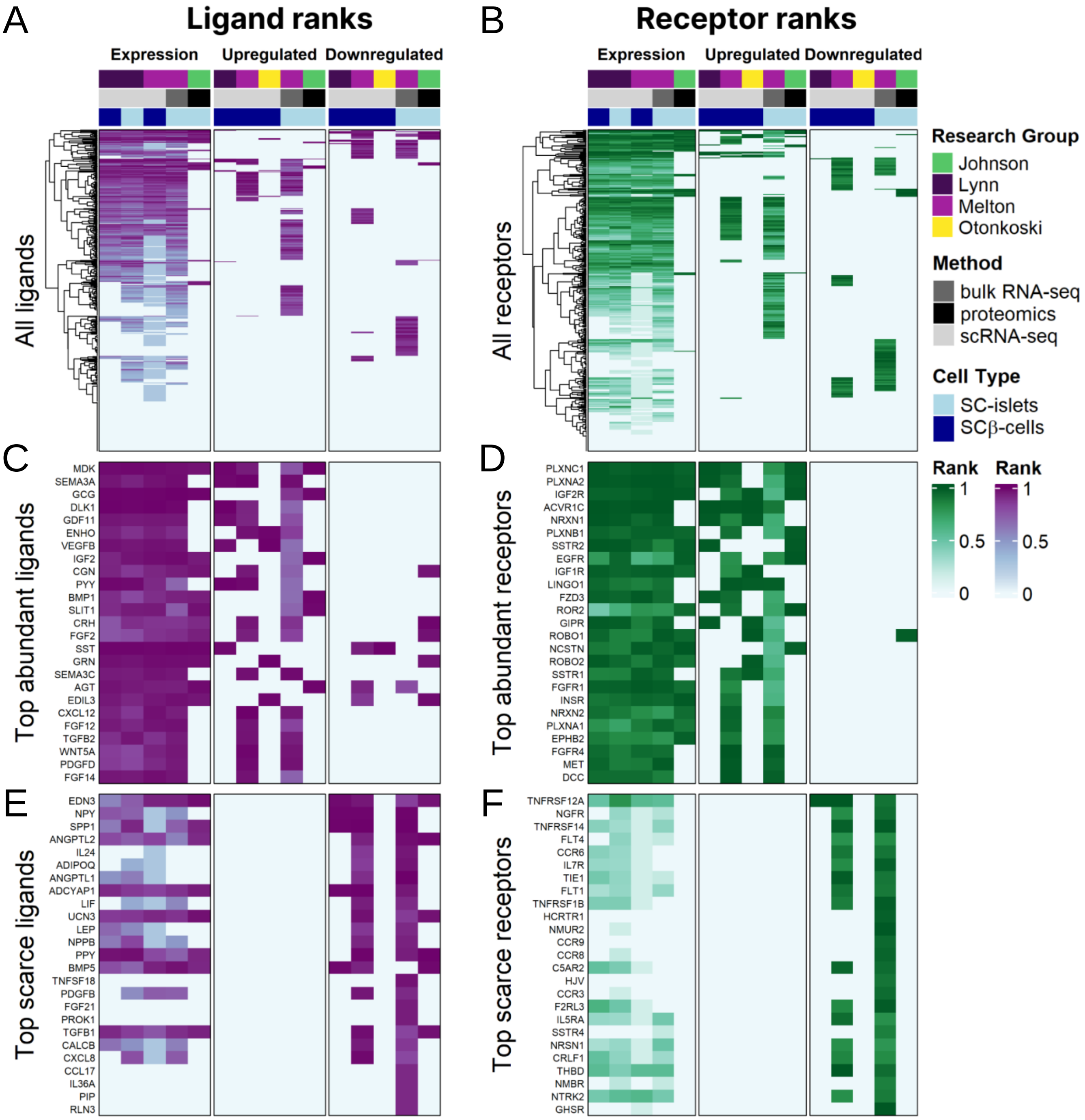


**Supplemental Figure 5. Ligand and receptor ranks in SC-islets based on 15 parameters. (A-F)** Ligand and receptor ranks across 15 parameters split into 3 categories: expression, upregulation, and downregulation. Each column corresponds to a parameter. Research group, cell type, and method for each parameter are indicated above the columns. **(A-B)** All 422 ligands **(A)** and 349 receptors **(B)**. **(C-D)** Top 25 abundant ligands **(C)** and receptors **(D)** in SC-islets. Abundant ligands and receptors are defined by high expression in SC-islets and higher expression in SC-islets compared to human islets. **(E-F)** Top 25 scarce ligands **(E)** and receptors **(F)** in SC-islets. Scarce ligands and receptors are defined by low expression in SC-islets and lower expression in SC-islets compared to human islets.

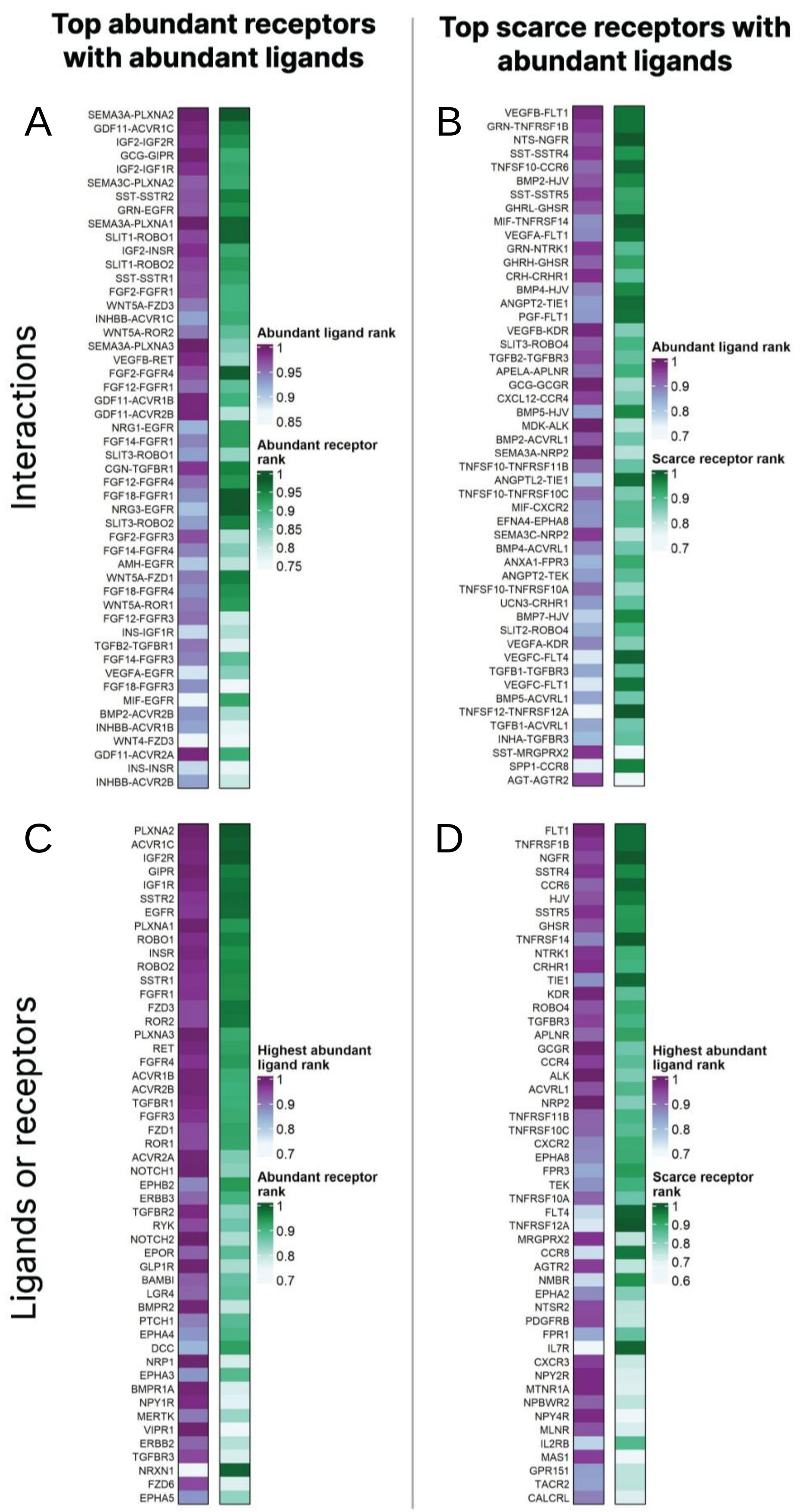


**Supplemental Figure 6. Prioritized ligand-receptor interactions in SC-islets.** Top 50 interactions in SC-islets between abundant receptors and abundant ligands **(A)**, scarce receptors and abundant ligands **(B)** **(C,D)** Top 50 abundant receptors with abundant ligands.

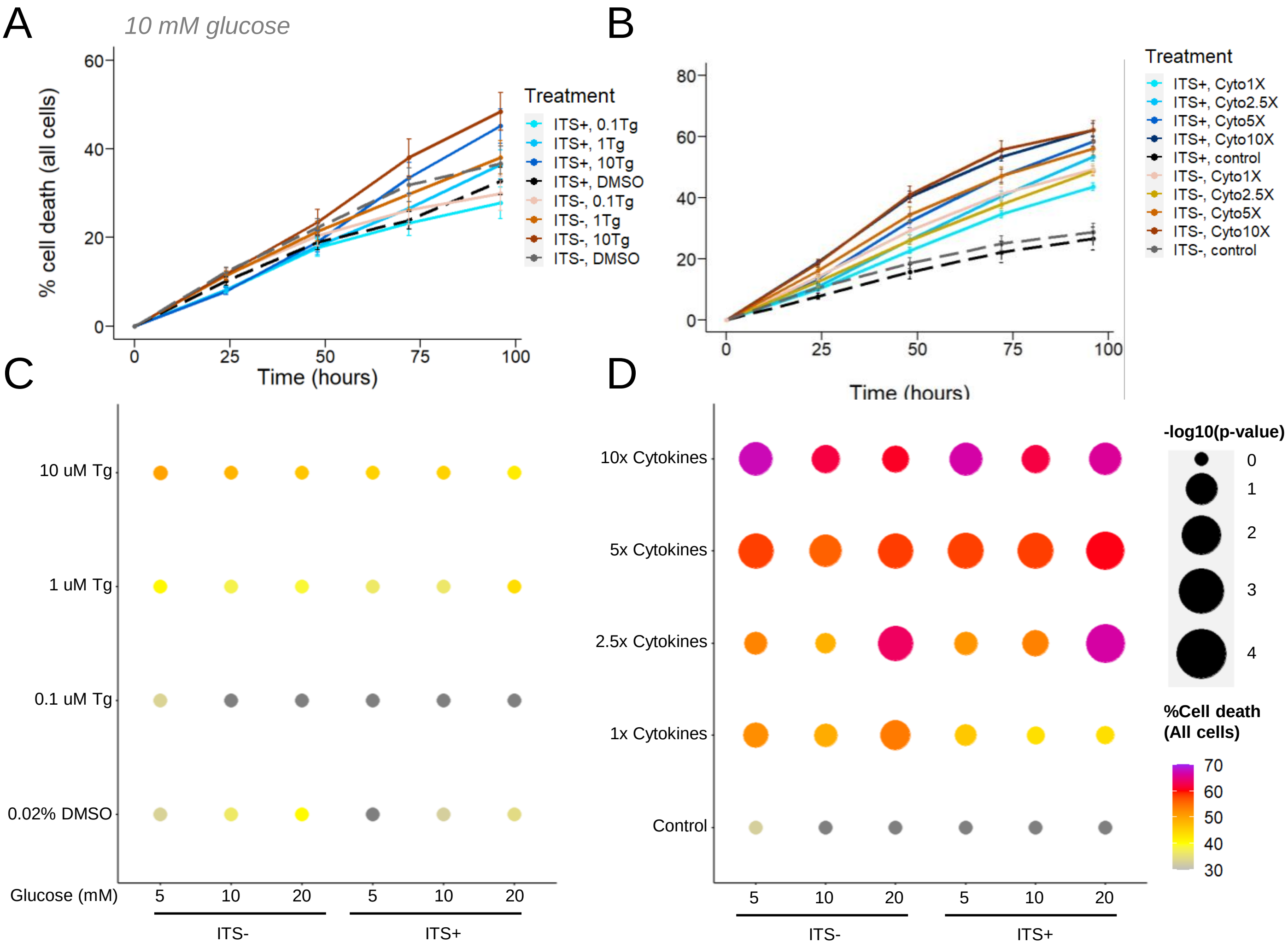


**Supplemental Figure 7. Stress assay on dispersed SC-islets using live cell imaging. (A)** Survival curves depicting dose dependent cell death induced by various concentrations of thapsigargin in media with 10 mM glucose. Error bars are SEM. **(B)** Survival curves depicting dose dependent cell death induced by various concentrations of cytokine cocktail at 10 mM glucose. Error bars are SEM. **(C)** Dot plot of total cell death thapsigargin conditions after 96 hours of imaging. Size of dots denotes –log10 (adj. p-value) compared to control. Color of the dots represent percentage of PI positive cells. **(D)** Dot plot of EGFP positive cell death in cytokine cocktail conditions after 96 hours of imaging. Size of dots denotes –log10 (adj. p-value) compared to control. Color of the dots represent percentage of PI positive cells. n=3 differentiations. Two-way ANOVA.

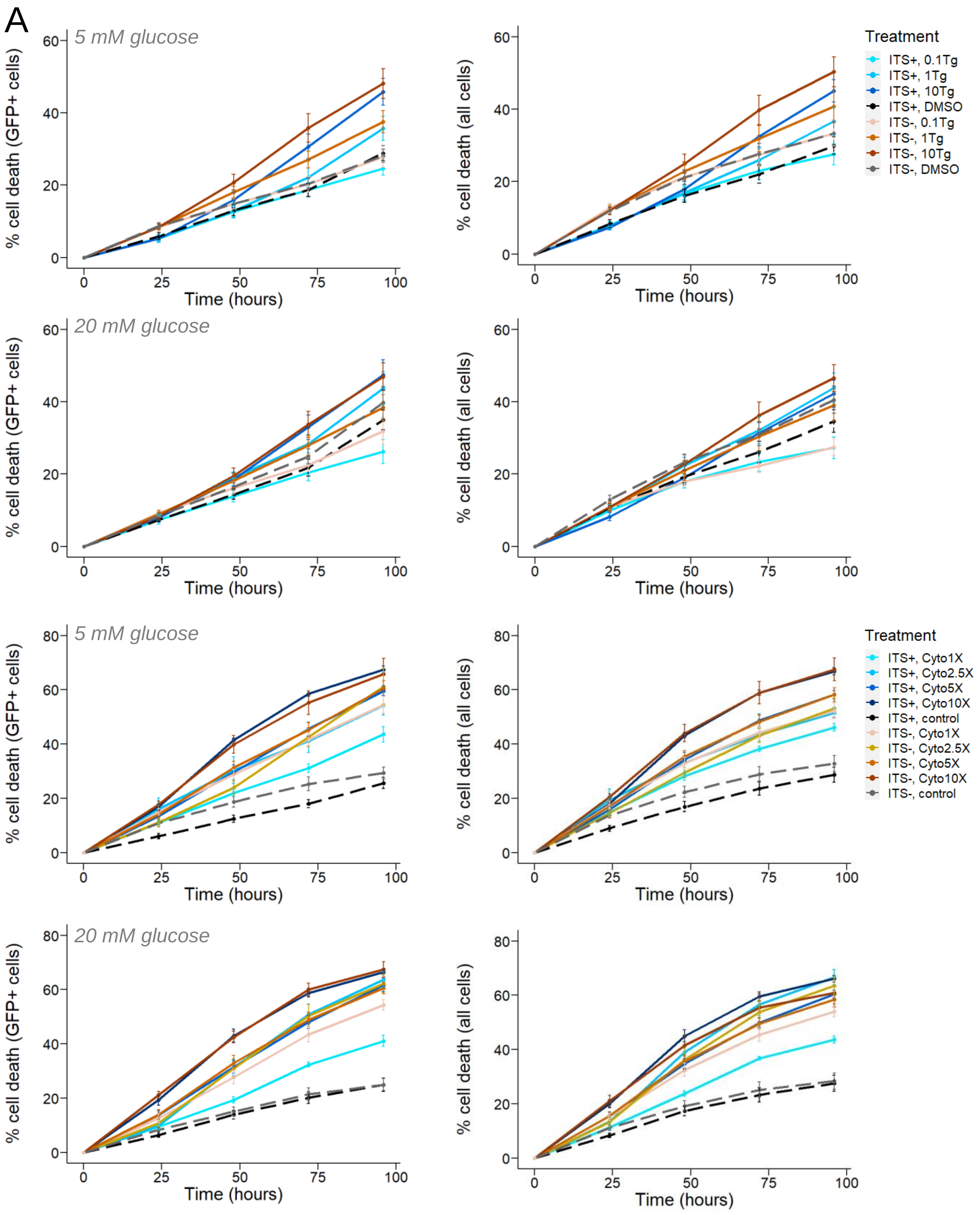


**Supplemental Figure 8. Stress assay on dispersed SC-islets using live cell imaging.** **(A)** Survival curves depicting dose dependent cell death induced by various concentrations of thapsigargin and cytokine cocktail at various glucose concentrations. Error bars are SEM. n=3 differentiations.

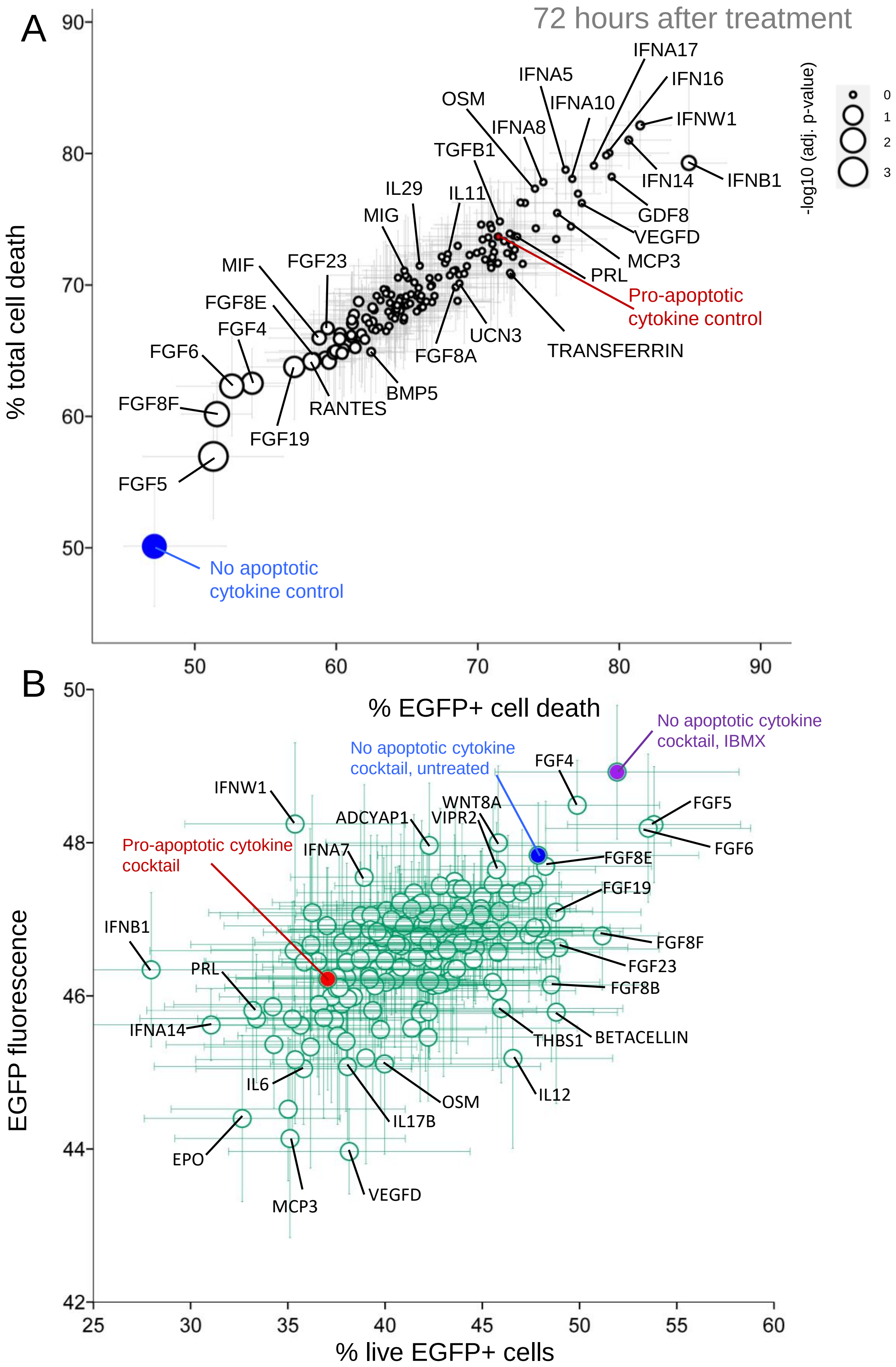


**Supplemental Figure 9. Cell count and EGFP fluorescence of the screen. (A)** Total cell death against EGFP positive cell death after 72 hours of treatment. Size of the dots denotes –log10 (adj. p-value) compared to pro-apoptotic cytokine control.**(B)** Average EGFP fluorescence in live cells against % live EGFP positive cells of all 148 ligands screened in the context of cytokine induction. n=5 differentiations.

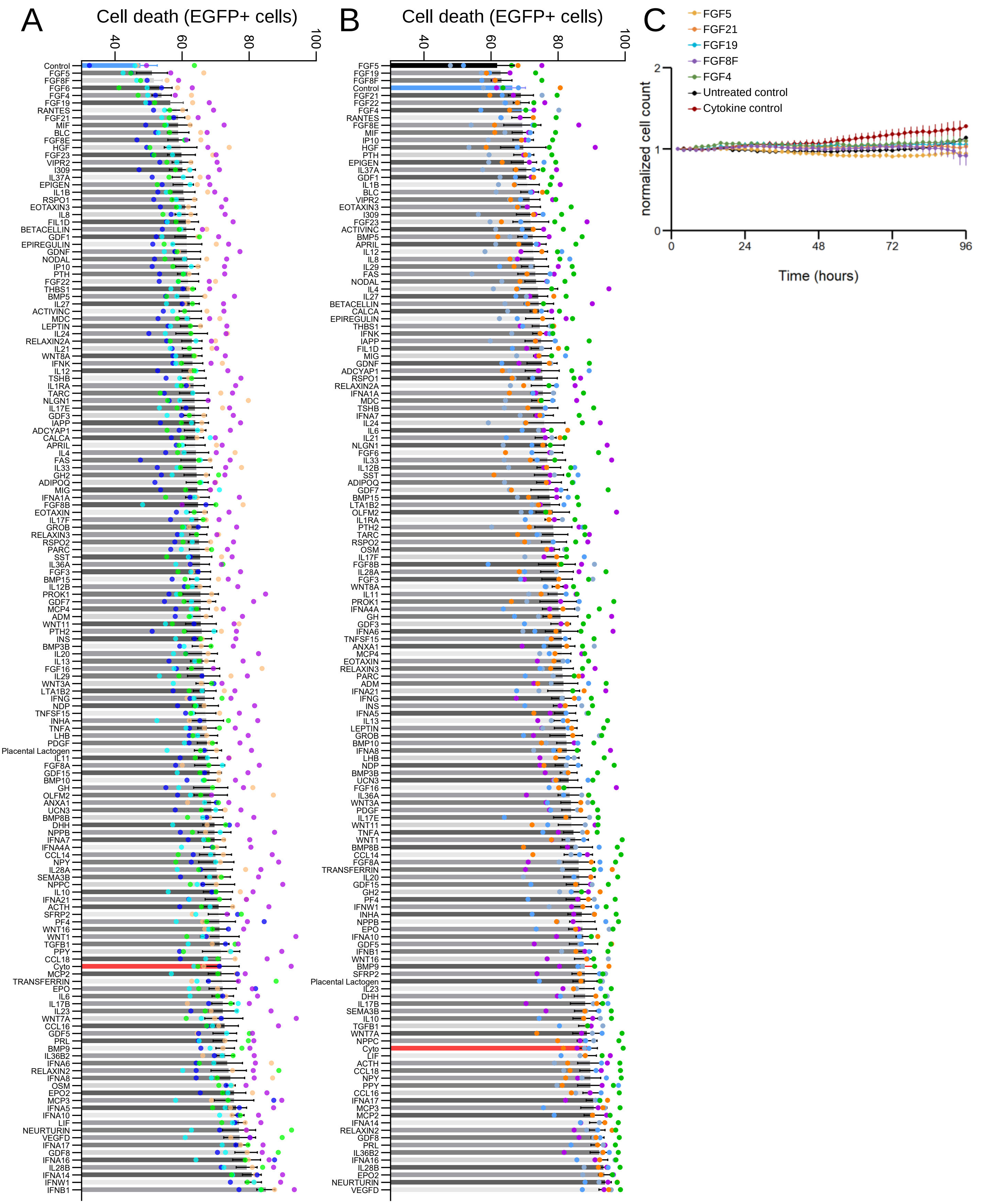


**Supplemental Figure 10. Screening for SCβ cell survival factors.** EGFP positive cell death screened at 72 hours **(A)** and at 96 hours **(B)**. Dots are biological replicate (n=5 differentiations). Blue bars are untreated controls, while red bars are cytokine treated pro-apoptotic controls. **(C)** Average cell count normalized to initial cell count in FGF trials. SEM error bars, * = adjusted p value <0.05. One-way ANOVA.

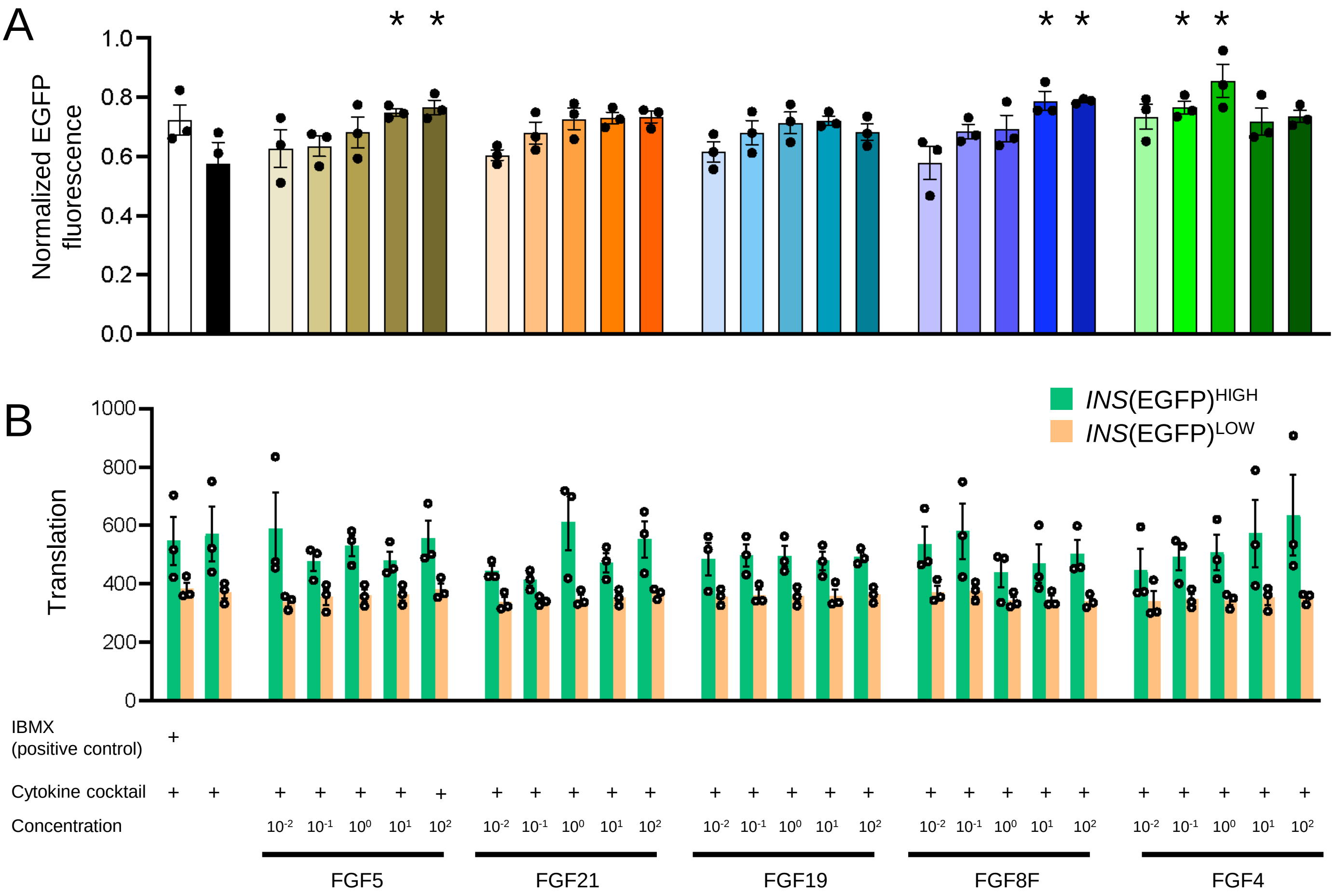


**Supplemental Figure 11. Dose response EGFP fluorescence and translational capacity of ligands of interest. (A)** Normalized EGFP fluorescence of ligands of interest. Fluorescence was normalized to the first 2 hours of imaging. n=3 differentiations. **(B)** Translational capacity in cells treated with ligands of interest, analyzed in the context of cells expressing high and low EGFP fluorescence. Error bars are SEM. Asterisks represent adjusted p value < 0.05 when compared with cytokine control. Two-way ANOVA. n=3 differentiations.
